# The Diversified Astrocyte Developmental Programs are Modulated by Primary Ciliary Signaling

**DOI:** 10.1101/2024.03.17.585433

**Authors:** Lizheng Wang, Qianqian Guo, Sandesh Acharya, Xiao Zheng, Vanessa Huynh, Brandon Whitmore, Askar Yimit, Mehr Malhotra, Siddharth Chatterji, Nicole Rosin, Elodie Labit, Colten Chipak, Kelsea Gorzo, Jordan Haidey, David Elliott, Tina Ram, Qingrun Zhang, Hedwich Kuipers, Grant Gordon, Jeff Biernaskie, Jiami Guo

## Abstract

Astrocyte diversity is greatly influenced by local environmental modulation. Here, we report that the vast majority of brain astrocytes across the entire brain possess a singular primary cilium, a specialized signaling antenna localized to cell soma. Comparative single-cell transcriptomics reveals that primary cilia mediate canonical Shh signaling to modulate astrocyte subtype-specific core features in synaptic regulation, intracellular transport, energy and metabolism. Independent of canonical Shh signaling, primary cilia are important regulators for astrocyte morphology and intracellular signaling balance. Dendritic spine analysis and transcriptomics reveal that perturbation of astrocytic cilia leads to disruption of neuronal development and global intercellular connectomes in the brain. Ultimately, mice with primary ciliary deficient astrocytes show behavioral deficits in sensorimotor function, sociability, learning and memory. Our results uncover a critical role for primary cilia in transmitting local cues that drive the region-specific diversification of astrocytes within the developing brain.

## Main

Cell function is constantly shaped through ever-changing interactions with the external environment. This is exemplified by astrocytes, an important class of glial cells in the brain. Astrocytes have emerged as highly versatile and critical regulators of nearly all aspects of brain development, homeostasis and plasticity; their diverse functional repertoire includes synaptic modulation, ion buffering, neurotransmitter uptake, integration within the blood-brain-barrier and immune reactivity^1–10^. During development, astrocytes throughout CNS establish communications with all other cell types (e.g., neurons, microglia, oligodendrocytes, pericytes, endothelial cells) and exhibit profound sensitivity to local cues in order to adopt unique functional attributes that are tuned to the demands within a given circuit or regional environment^11–22^. While we have begun to appreciate astrocyte heterogeneity in the brain, little is known about how different subtypes of astrocytes are created during development to differentially support their local neural circuits. This gap of knowledge is critical, considering astrocyte dysfunction and their diverse responses to pathological conditions are major components underlying a wide range of neurodevelopmental and neurological disorders^5, 23–28^.

The fact that astrocytes show diverse features not only specific to brain regions but also within the same region indicates that differences both in the local environment and how astrocyte subtypes interpret environmental cues are important for their heterogeneity specification^11–22^. To study how this environment – genetic interaction is integrated to specify astrocyte diversity, we focused on primary cilia, small signaling antennae protruding from the cell soma. Housing a concentrated, diverse cast of signaling components [e.g., G protein-coupled receptors (GPCRs), receptor tyrosine kinases, ion channels, Ca^2+^] to detect extracellular cues^29–35^, primary cilia are increasingly recognized as underappreciated signaling hubs indispensable for coordinating major cellular functions in response to extracellular environment. The significance of primary cilia in the brain is evidenced in ciliopathies^34, 36–42^, referring to 30+ human disorders caused by genetic mutations of ciliary genes, where dysfunctional cilia lead to brain structural defects and functional deficits associated with neurodevelopmental disorders such as intellectual disabilities (ID), autism spectrum disorders (ASD), and epilepsy^43–50^. Nevertheless, our mechanistic understanding of how cilia function in the brain is rudimentary. We and others have recently demonstrated that primary cilia of neurons are critical for regulating all major neuronal developmental events ranging from neurogenesis, migration, morphogenesis, axonal development, and synapse formation^34, 40, 41, 51–57^. Signaling pathways such as Shh, Wnt, Notch, GPCR, PDGF, and FGF that are known to modulate astroglial fate specification and function have been reported to be mediated through primary cilia^17, 58–60^. However, the role of primary cilia in astrocytes remains largely unexplored.

These findings prompted us to ask several key questions. First, do all subtypes of astrocytes possess primary cilia and if so, do they rely on primary cilia to transduce signaling events relevant for shaping astrocyte subtype heterogeneity and maturation? Second, considering the severe brain deficits associated with ciliopathies and the importance of astrocytes on neuronal development, what are the functional consequences of cilia-deficient astrocytes on neuronal development and behaviors?

By combining mouse genetics, comparative single-cell transcriptomics, and mouse behavioural analyses, we analyzed how primary ciliary signaling modulate astrocyte subtype-specific maturation and assessed the impact of ciliary deficient astrocytes on neuronal programs and behaviors. Our data uncovered a surprisingly important role for primary cilia in astrocyte development, by acting as key sensors of local environmental cues that drive astrocyte maturation and regional-specific functional specification to support brain development.

## Results

### Almost all astrocytes possess a singular primary cilium in the brain

We first examined if all astrocytes possess primary cilia in the brain. To label individual astrocytes, we used the *Aldh1l1-Cre^ERT^*^2^ mouse line that drives Cre expression specifically in all astrocytes in a tamoxifen inducible manner. The Ai9 reporter was bred in to generate the *Aldh1l1-Cre^ERT2^; Ai9* mouse line, which allows permanent labeling of developing and adult astrocytes with tdTomato (tdTom) upon tamoxifen-driven Cre recombination. Because astrocytes show heterogeneity across different brain regions and within the same region, we focused our analysis on the cerebral cortex (including the grey matter protoplasmic astrocytes in the upper layers II/III, deeper layers V/VI, corpus collosum where fibrous astrocytes reside, striatum (dorsal), hippocampus (CA1), thalamus, hypothalamus, and the cerebellum (including the Bergmann glia and the velate astrocytes) (Fig. 1a). Primary cilia were detected by immunofluorescence staining for Arl13b protein that labels all primary cilia^45^. With the exception of the fibrous astrocytes, the vast majority of astrocytes, regardless of region or subtype, possess a singular primary cilium (Fig. 1b). Interestingly, only 38.02 ± 3.28 % of astrocytes in the white matter possess primary cilia labeled with anti-Arl13b. To rule out the possibility of lower (or undetectable) Arl13b expression levels in the fibrous astrocytes, we further validated this result using a transgenic mouse over-expressing Arl13b-mCherry and similarly found only 34.51 ± 3.00% of fibrous astrocytes were ciliated (Fig.1b). Interestingly, astrocyte primary cilia exhibited a regional variation in ciliary length: the deeper cortical protoplasmic (3.78 ± 0.07 µm) and CA1 hippocampal astrocytes have the longest ciliary length (3.80 ± 0.07 µm), whereas the cerebellar Bergmann glia have the shortest cilia (2.51 ± 0.07 µm) (Fig. 1c, d).

**Fig. 1:**
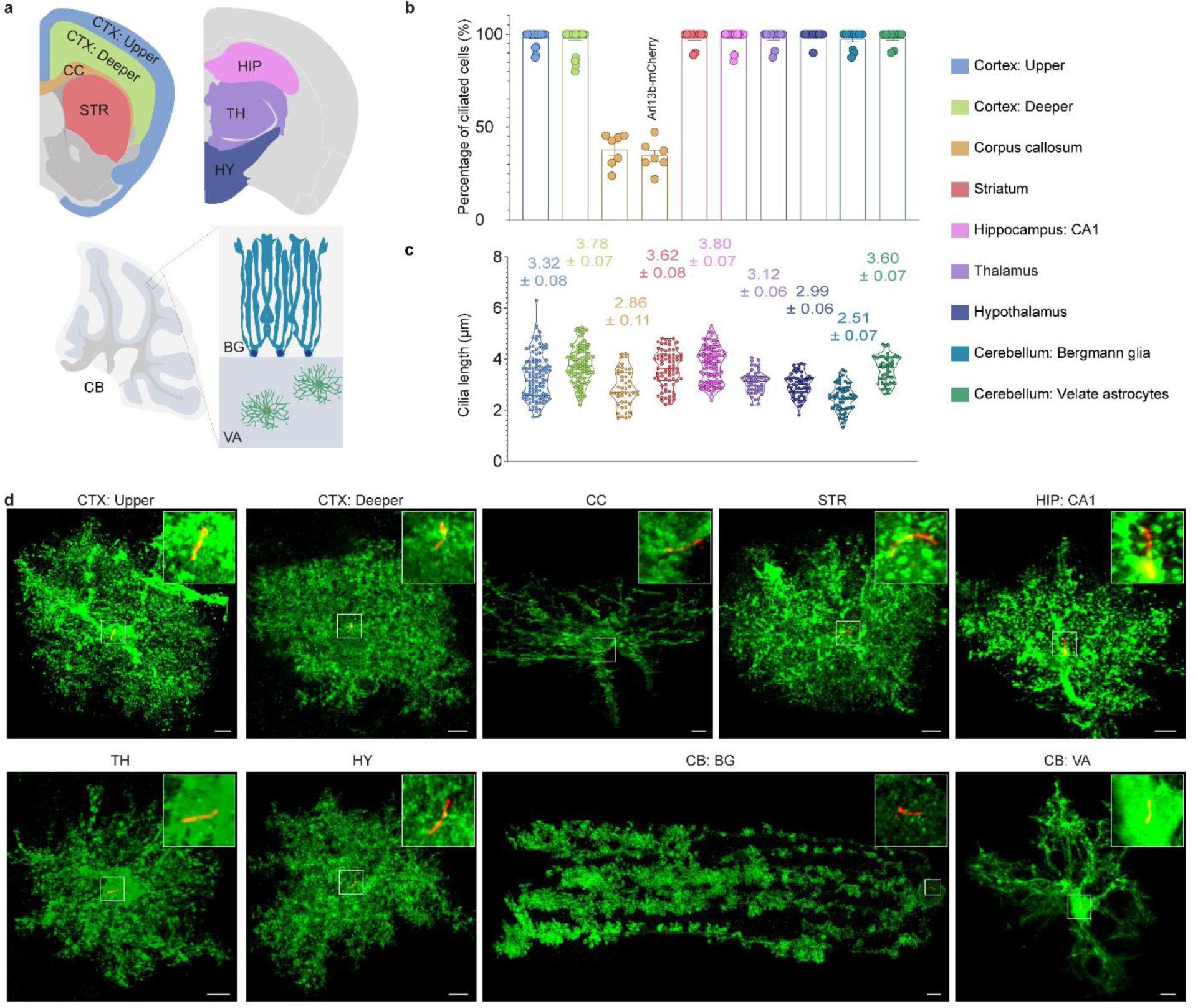
Primary cilia are present in the majority of astrocytes. **(a)** Illustration of brain regions and astrocyte subtypes for primary cilia analysis. CTX, cortex; CC, corpus callosum; STR, striatum; HIP, hippocampus; TH, thalamus; HY, hypothalamus; CB, cerebellum; BG, Bergmann glia; VA, velate astrocytes. **(b, c)** Percentage of ciliated cells and the length of primary cilia in astrocytes. “Arl13b-mCherry” in (b), the Arl13b-mCherry transgenic mouse line. Data are shown as means ± SEM. N= 7-32 images per brain region in (b), 46-110 cilia per brain region in (c). **(d)** Representative images of primary cilia in different astrocyte subsets. AAV5-GFAP-mCherry was injected to the lateral ventricle at P0 to label astrocyte morphology, and brains were collected at P60. Primary cilia were labeled with anti-Arl13b antibody. Green, mCherry; Red, Arl13b. Scale bars, 5 µm.

### Primary ciliary signaling differentially modulate astrocyte subtype-specific transcriptional signatures

The presence of primary cilia on distinct astrocyte subtypes prompted us to ask if astrocytes utilize primary cilia to detect environmental signals to help configure their diversified developmental programs. To answer this, we generated two astrocyte specific ciliary mutant mice, using a conditional mouse *Arl13b^lox/lox^* allele of *ARL13B*^45, 61^, and the *Ift88^lox/lox^* allele of *IFT88*^62, 63^, respectively, bred with *Aldh1l1-Cre^ERT2^* (*Arl13b^cKO^* and *Ift88^cKO^*). *ARL13B* is a ciliopathy causative gene that encodes a cilia-specific small GTPase critical for ciliary signaling (e.g. Shh, cilia-GPCR signaling)^34, 41, 42, 64^. Deficiency of Arl13b results in malformation of primary cilia with defective Shh and cilia-GPCR signaling^34, 41, 42, 64^. *Ift88* encodes a cilia-specific protein essential for ciliary formation, and Ift88 deficiency results in a complete ablation of the primary ciliary structure^65^. The Cre expression was induced by tamoxifen injections at P7-9, bypassing astrogliogenesis in the first postnatal week and before the week 4 when astrocytes reach morphological maturity^15, 66^. We confirmed that tamoxifen mediated highly efficient Cre recombination in around 93% of all astrocytes (S100β^+^) (Extended Data Fig. 1a, b) and complete loss of primary cilia in all Cre^+^ *Ift88^cKO^* astrocytes *in vivo* (Extended Data Fig. 1c). Using acetylated tubulin as a ciliary marker^67^, we also confirmed that *Arl13b^cKO^* astrocytes primary cilia are significantly shorter than control (Extended Data Fig. 1d, e), consistent with the known effects of Ift88 or Arl13b deficiency on primary cilia formation^45, 65, 68^

Single-cell RNA sequencing (scRNAseq) (Extended Data Fig. 2a-d) was performed on astrocytes isolated from the cerebral cortices (including the hippocampus) from *Arl13b^cKO^*, *Ift88^cKO^*, and control mice, and the cerebellum from *Arl13b^cKO^* and control mice at postnatal day 56 (P56). In the cerebral cortex, cortical white matter, and the hippocampus, we identified 5 distinct astrocyte subtypes (Fig. 2a) that can be partially identified by layer- and region-specific markers. Cluster 0, 1, 2 express grey matter genes *Nupr1* and *Thrsp*, representing three major protoplasmic cortical astrocytes^12^. Specifically, Cluster 0 is enriched with upper cortical layer markers *Mfge8* and *Igfbp2*, while Cluster 1 and Cluster 2 show mixed expression of both upper (*Mfge8*, *Igfbp2*) and lower markers (*Nnat*, *Agt*). Cluster 3 (Gfap^high^) expresses white matter astrocyte genes *Vim* and *Gfap*, representing the fibrous astrocytes. Cluster 4 (Crym^+^) are enriched for *Prss56* and *Crym* genes reported to be enriched in subventricular zone (SVZ) and neural stem cells (Extended Data Fig. 2e).

**Fig. 2:**
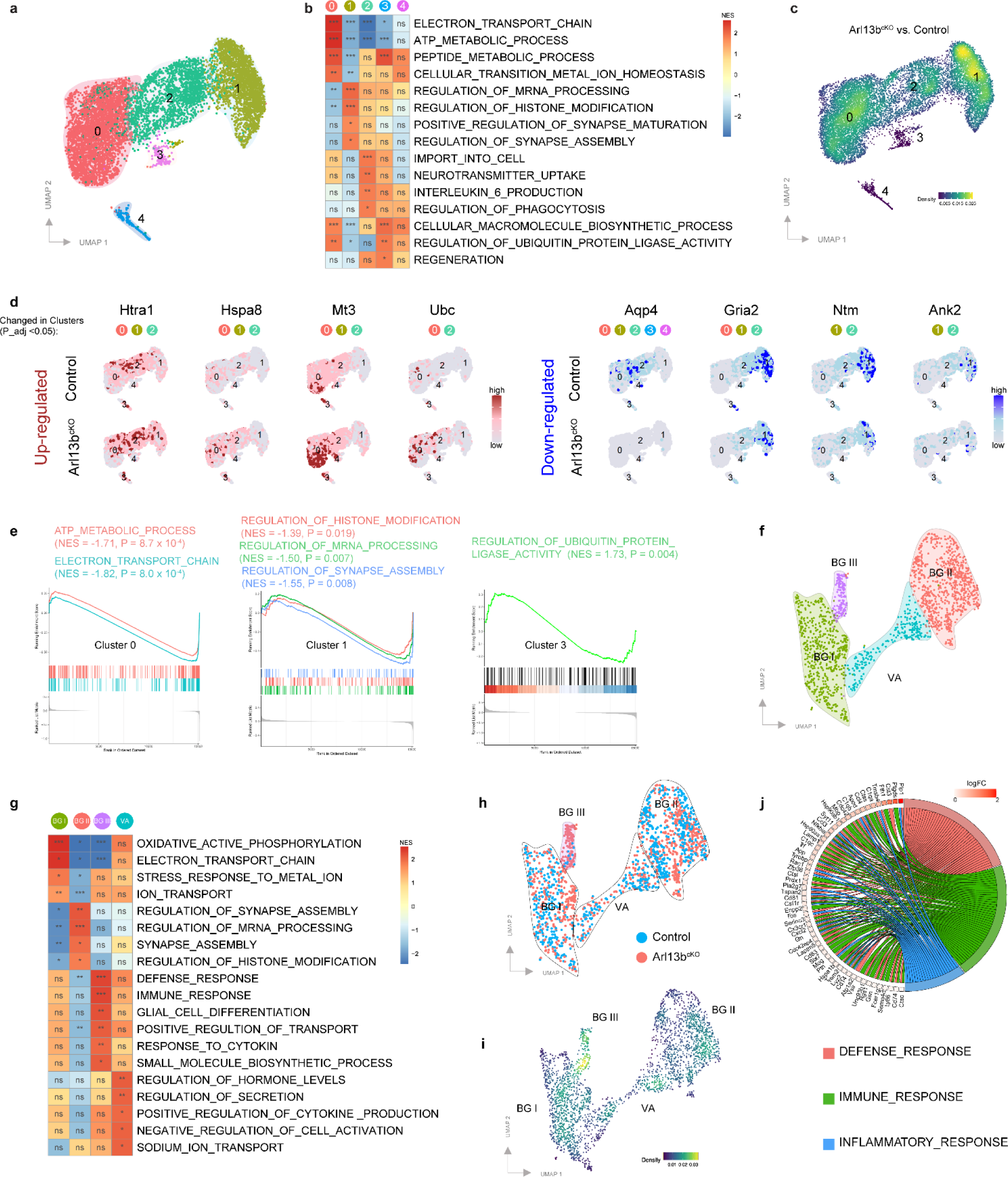
Single-cell transcriptomics analysis of *Arl13b^cKO^* astrocytes. **(a)** Uniform manifold approximation and projection (UMAP) plot of cortical astrocytes (including hippocampus and corpus callosum) from *Aldh1l1Cre^ERT2^* (Control) and *Arl13b^lox/lox^; Aldh1l1Cre^ERT2^* (*Arl13b^cKO^*) mice. **(b)** Enriched GO biological processes identified by GSEA analysis in cortical astrocyte clusters. NES, normalized enrichment score. **(c)** Kernel density estimates depicting magnitude of molecular response elicited by *Arl13b* deletion, calculated by summing DEG FCs (*Arl13b^cKO^* vs. Control) for each cortical astrocyte cluster. **(d)** Visualization of representative up- and down-regulated DEGs in *Arl13b^cKO^* cortical astrocyte clusters. **(e)** The influence of *Arl13b* deletion on the key GO pathways in cortical astrocyte clusters. **(f)** UMAP plot of cerebellar astrocytes from control and *Arl13b^cKO^* mice. **(g)** Enriched GO biological processes identified by GSEA analysis in cerebellar astrocyte clusters. **(h)** UMAP plot colored by sample groups. **(i)** Magnitude of molecular response in *Arl13b^cKO^* cerebellar astrocytes, calculated by summing DEG FCs (*Arl13b^cKO^* vs. Control) for each cerebellar astrocyte cluster. **(j)** Circos plot of enriched GO terms in BGIII.

Gene set enrichment analysis (GSEA) associated with biological processes revealed diverse functional features in these astrocyte subpopulations (Fig. 2b, Extended Data Fig. 2f, g): Cluster 0 is associated with electron transport chain, ATP metabolism, peptide metabolism, and ion homeostasis; Cluster 1 is associated with synapse maturation/assembly, RNA processing, and histone modification; Cluster 2 is associated with cellular transport, neurotransmitter uptake, cytokine production, and phagocytosis; Cluster 3 shows enriched pathways with macromolecule biosynthesis, ubiquitin ligase activity, and regeneration. Based on these pathway analyses, we annotated Cluster 1 as being specifically associated with “synaptic regulation”, while Clusters 0 and 2 were associated with “metabolism and homeostasis regulation”.

We next interrogated how primary cilia influenced astrocyte subtype-specific expression profiles. In *Arl13b^cK^*^O^ and *Ift88^cKO^* brains, although all astrocyte subpopulations were identified and no apparent difference in subtype composition was found compared to controls (Extended Data Fig. 5m), the global magnitude of differential expressed genes (DEGs) revealed marked perturbations in *Arl13b^cKO^* and *Ift88^cKO^* all 3 cortical clusters (Fig. 2c, Extended Data Fig. 3a, b). Only 1 DEG (*Aqp4*, downregulated) was found in Cluster 3 in *Arl13b^cKO^* (0 in *Ift88^cKO^*) and 1 DEG in Cluster4 in *Arl13b^cKO^* (6 in *Ift88^cKO^*). Among Clusters 0-2, top DEGs in *Arl13b^cKO^* are similarly changed in *Ift88^cKO^* and differentially distributed to different clusters (Fig. 2d, Extended Data Fig. 3c). Importantly, although we found more DEGs in *Ift88^cKO^* clusters, GSEA analysis revealed that the top enriched pathways in *Arl13b^cKO^* were similarly altered in *Ift88^cKO^* clusters, suggesting that the DEGs induced by the two independent ciliary gene deletion converge on the same signaling pathways and cellular processes in astrocytes (Extended Data Fig. 3d-f). GSEA analysis also revealed a conserved enrichment in response to ER stress/protein folding (Cluster 0, 1, 2 in *Ift88^cKO^*), amyloid precursor protein metabolic process (Cluster 0, 1, 2 in *Ift88^cKO^*), and GPCR signaling pathway (Cluster 0 in *Ift88^cKO^*) across Cluster 0, 1, 2 in *Arl13b^cKO^*. However, more pathways are differentially disturbed among different clusters in *Arl13b^cKO^* and *Ift88^cKO^* astrocytes (Extended Data Fig. 3d-f). For example, *Arl13b^cKO^* and *Ift88^cKO^* Cluster 0 shows enrichment in PI3K signaling and glial cell differentiation but downregulated ATP production pathways (Extended Data Fig. 3d); Cluster 1 shows enrichment in response to extracellular stimulus but downregulated synapse assembly (Extended Data Fig. 3e); Cluster 2 shows upregulated response to growth factor stimulus but downregulated transmembrane transport and ion homeostasis pathways (Extended Data Fig. 3f).

*Arl13b^cKO^* and *Ift88^cKO^* astrocytes also show significant changes in the key gene signatures that distinguish each cluster: *Arl13b^cKO^* and *Ift88^cKO^* Cluster 0 show attenuated gene signature for ATP metabolism and electron transport chain; *Arl13b^cKO^* and *Ift88^cKO^* Cluster 1 show attenuated gene signature for RNA processing, histone modification, and synapse assembly; While in Cluster 3, *Arl13b^cKO^* astrocytes show elevated ubiquitin ligase activity pathways compared to controls (Fig. 2e, Extended fig. 3g).

In the cerebellum, we identified 4 astrocyte subtypes, including three BG clusters and one VA cluster (Fig. 2f, Extended Data Fig. 4a). Interestingly, BGI and BGII share similar enriched pathways with cortical astrocyte Clusters 0 and 1, respectively, whereas the VA cluster shows similarity with cortical Cluster 2 (Fig. 2g, Extended Data Fig. 4b). Similar as cortical astrocytes, gene signatures of *Arl13b^cKO^* CB astrocytes are differentially disturbed among different clusters in pathways related to synapse regulation, ATP production, transmembrane transport, water/ion homeostasis, MAPK signaling, and immune response/cytokine production (Extended Data Fig. 4c-f). Notably, BGIII cluster consisted almost entirely of *Arl13b^cKO^* astrocytes (Fig. 2h), suggesting that primary cilia dysfunction results in the emergence of a novel BG subtype. Indeed, the BG III cluster exhibited the greatest degree of transcriptomic perturbation following cilia dysfunction (Fig. 2i) that included an enrichment of genes associated with immune response, defense response, and response to cytokines, all indicative of a reactive astrocyte signature (Fig. 2g, j). Interestingly, compared to known reactive astrocyte markers, including the Pan-reactive, A1-like, A2-like, and Alzheimer’s disease associated astrocyte markers^69^, BGIII specifically showed an upregulation in A2-like astrocyte marker genes and no alteration in the others (Extended Data Fig. 4g), indicating that ciliary dysfunction severely altered the gene expression of a BG subset specifically towards the A2-like reactive astrocytes at the resting stage.

Taken together, these results demonstrate that the two ciliary mutants show similar transcriptional alterations that converge onto the same signaling pathways in astrocytes, suggesting that dysfunction of primary cilia markedly alters astrocyte transcriptional programs in the cerebral cortex and the cerebellum, resulting in substantial subtype-specific changes in astrocyte development and function.

### Primary cilia modulate astrocyte transcriptomes partly by canonical Shh signaling pathway

Prior studies demonstrated that layer V neurons in the cerebral cortex and Purkinje cells in the cerebellum secrete Shh that acts to regionally specify and maintain astrocyte diversity^11, 18, 19, 21, 70^. Primary cilia serve as a primary mediator for canonical Shh (cShh) signaling transduction in most mammalian cell types but their role in mediating cShh signaling in astrocytes has not been examined^29, 71, 72^. The translocation of Smo into primary cilia upon Shh stimulation is a hallmark of cShh signal transduction^73, 74^. We found that Smo showed enriched localization at primary cilia in cultured cortical astrocytes in response to Shh stimulation (Fig. 3a-c). Further, using a *SmoM2*^75^ allele (*SmoM2-eYFP*), which expresses the constitutively active form of Smo in a Cre-dependent manner, we observed over 82% of astrocytes show ciliary localized SmoM2 *in vivo* (Extended Data Fig. 5a, b), suggesting that primary cilia transduce cShh signaling in astrocytes.

**Fig. 3:**
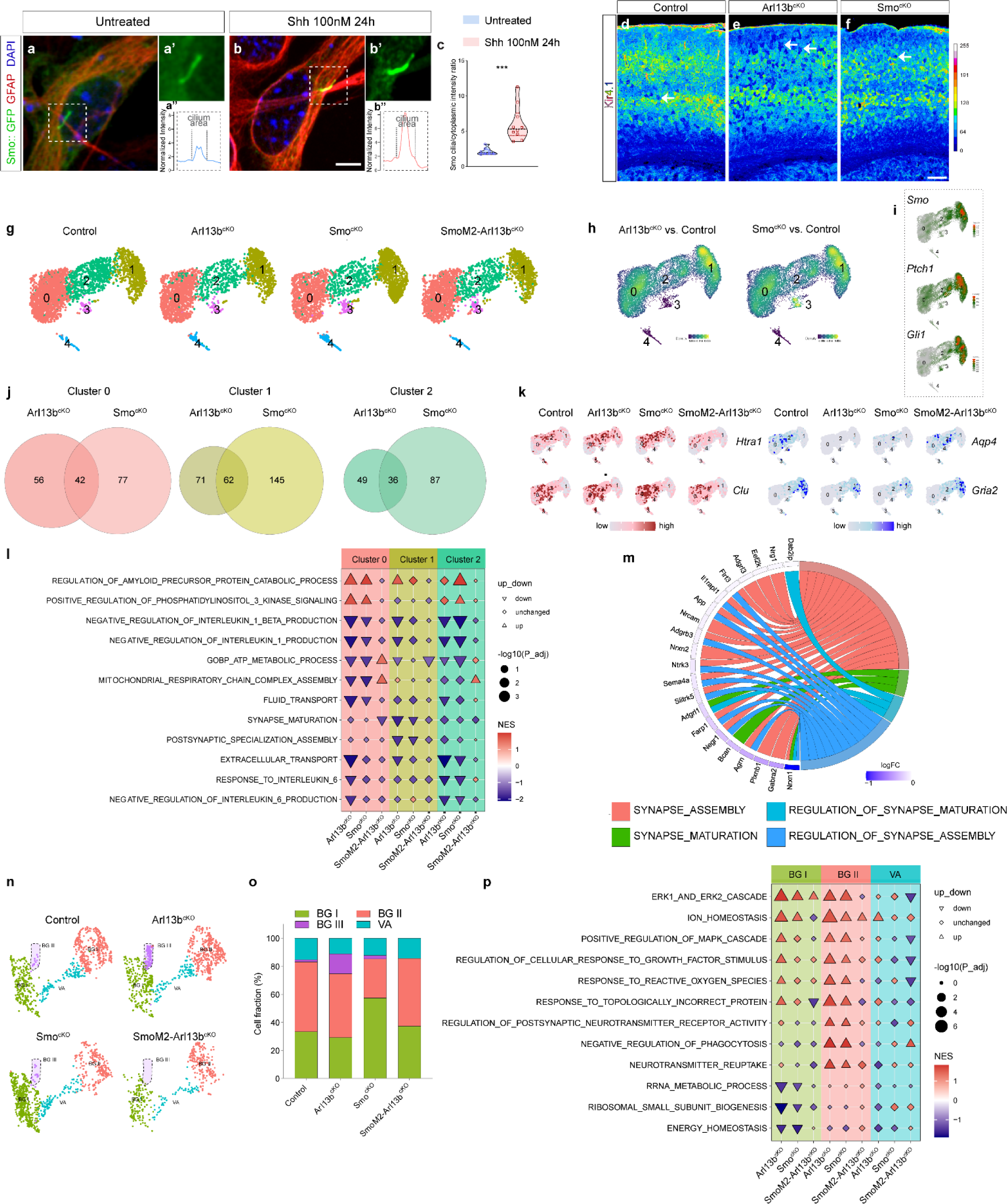
Primary cilia modulate astrocyte transcriptomes partly via canonical Shh signaling pathway. **(a-c)** The ciliary enrichment of Smo in cultured mouse astrocytes upon SHH treatment. Mouse cerebral cortical astrocytes were cultured and transfected with Smo::GFP plasmid. Cells were treated with SHH at 100nM for 24h post transfection. **(a’, b’)** Enlarged images of the boxed area in A, B. Scale bar, 5 µm. **(a’’, b’’)** Representative Smo intensity profiles of ciliary and cytoplasmic area. Unpaired t-test was employed to analyze the differences. *\*\*\*P* < 0.001.**(c)** Quantification of Smo cilia/cytoplasmic intensity ratio. N= 8-10 astrocytes per group. **(d-f)** Kir4.1 expression pattern in control, *Arl13b^cKO^* and *Smo^cKO^* mouse cortex shown pseudocolored for intensity. Scale bar, 200 µm **(g)** UMAP plots of cortical astrocytes from control, *Arl13b^cKO^, Smo^cKO^* and *SmoM2-Arl13b^cKO^* mice. **(h)** Kernel density estimates depicting magnitude of molecular response in *Arl13b^cKO^* and *Smo^cKO^*, calculated by summing DEG FCs for each cortical astrocyte cluster. **(i)** Expression patterns of canonical Shh signaling target genes in cortical astrocyte clusters. **(j)** Venn diagrams of DEGs found in *Arl13b^cKO^* and *Smo^cKO^* groups (compared to Control) across cortical astrocyte Clusters 0, 1, 2. **(k)** Expression of representative genes regulated by Cilia-cShh signaling in cortical astrocytes. **(l)** Top GO biological processes regulated by Cilia-cShh signaling in cortical astrocytes. **(m)** Circos plot of representative “synapse” associated GO terms downregulated in *Arl13b^cKO^* Cluster1 cortical astrocytes. **(n)** UMAP plots of cerebellar astrocytes from Control, *Arl13b^cKO^, Smo^cKO^* and *SmoM2-Arl13b^cKO^* mice. **(o)** Cell percentage of cerebellar astrocyte subtypes in each sample group. **(p)** GO biological processes regulated by Cilia-cShh signaling in cerebellar astrocytes.

We next asked if primary cilia mediate cShh signaling to influence astrocyte gene expression. Kir4.1 (potassium inwardly rectifying channel) and GluA1 (glutamate ionotropic receptor AMPA type subunit 1) are astrocyte-specific proteins important for their function and known to be directly regulated by Shh signaling^11, 19, 76^. In controls, Kir4.1 is highly expressed in layer II/III and layer V cortical protoplasmic astrocytes, hippocampal astrocytes, and BGs, but is expressed at low level in VAs (Fig. 3d, Extended Data Fig. 5c, e, f). In *Arl13b^cKO^* (*Arl13b^lox/lox^; Aldh1l1-Cre^ERT2^*) and *Ift88^cKO^* (*Ift88^lox/lox^; Aldh1l1-Cre^ERT2^*) brains, we found a “patchy” pattern of Kir4.1 expression, with a strong reduction of Kir4.1 expression in subsets of cortical protoplasmic astrocytes across the cortex (Fig. 3e, Extended Data Fig. 5d), as well as the hippocampus and the cerebellum (Extended Data Fig. 5e, f). A very similar pattern but to a less extent was observed in *Smo^cKO^* (*Smo^lox/lox^; Aldh1l1-Cre^ERT2^*) brain, where Shh loss-of-function is specifically induced in all astrocytes (Fig. 3f, Extended Data Fig. 5e, f). Interestingly, in the *Arl13b-hGFAP-Cre^cKO^* (*Arl13^blox/lox^; hGFAP-Cre*) cerebellum, in which hGFAP-Cre leads to gene deletion in astroglial progenitors^77^, we found not only a reduction of Kir4.1 expression in subsets of mutant BGs, but also elevated expression in adjacent VAs (Extended Data Fig. 5e), indicating deficiency of primary ciliary during the progenitor stage expanded the impact on astrocyte subtype-specification. Notably, we also observed this elevated expression of Kir4.1 in VAs in the *Ift88^cKO^* cerebellum (Extended Data Fig. 5e), suggesting that complete ciliary ablation induced in *Ift88^cKO^*, compared to the partial ciliary defects induced in *Arl13b^cKO^*, induced a more severe impact on astrocyte subtype-specification. In controls, GluA1 is highly expressed in BGs and hippocampal astrocytes, but low in VAs and protoplasmic astrocytes across the cerebral cortex (Extended Data Fig.5g, h). We found that both *Arl13b^cKO^* and *Ift88^cKO^, as well as Smo^cKO^* brains showed reduced expression of GluA1 in the hippocampus and the cerebellum (Extended Data Fig.5g, h). Together, these results suggest that primary cilia mediate cShh to regulate astrocyte gene expression.

To further assess the impact of Cilia-cShh signaling on astrocyte transcriptomes, we performed scRNAseq analysis comparing control mice with *Arl13b^cKO^, Ift88^cKO^, Smo^cKO^,* and *SmoM2-Arl13b^cKO^*. *SmoM2-Arl13b^cKO^* are the Shh “Gain-of-function (GOF)” mice in which SmoM2 is expressed in *Arl13b^cKO^* astrocytes to engage downstream Shh signaling activation even in the absence of functional cilia. We confirmed that *SmoM2* is expressed in ∼98% of Cre^+^ astrocytes and that the SmoM2^+^ *Arl13b^cKO^* astrocytes showed complete loss of Arl13b protein (Extended Data Fig. 5i-l). We reasoned that if primary cilia modulate astrocyte transcription through Shh signaling, changes in *Arl13b^cKO^* would overlap with *Smo^cKO^* mice and be largely corrected in *SmoM2-Arl13b^cKO^* mice. The differences between *Arl13b^cKO^* and *Smo^cKO^* mice would shed light on the impact of primary cilia mediated cShh independent signaling on astrocyte transcriptomes.

Global transcriptomic analysis showed shared transcriptomic profiles in *Arl13b^cKO^* and *Smo^cKO^* astrocytes (Fig. 3g, h, Extended Data Fig. 5m), with the most pronounced gene expression changes in Cluster 1, followed by Cluster 2, which correlates with Shh signaling activity level (Fig. 3i). Assessment of DEGs shared between *Arl13b^cKO^* and *Smo^cKO^* groups highlighted a molecular overlap that constitutes 42.9% (42/98 genes) of DEGs in Cluster 0, 46.6% (62/133 genes) in Cluster 1, and 42.4% (36/85 genes) in Cluster 2 in *Arl13b^cKO^* (Fig. 3j).

To further define the primary cilia-mediated cShh signaling gene signature (“Cilia-cShh”) in astrocytes, we selected genes that show similar changes in both *Arl13b^cKO^* and *Smo^cKO^* and that are “rescued” in *SmoM2-Arl13b^cKO^* compared to controls. This identified 50 down regulated “Cilia-cShh” genes in all clusters combined, such as *Aqp4* (Aquaporin 4, down in all clusters), which encodes the predominant water channel in the brain critical for water homeostasis^78^ and *Gria2* (down in cluster 1 and 2) that encodes a AMPA receptor subunit 2 that regulate astrocyte activity^8^ (Fig. 3k). 26 “Cilia-cShh” genes were upregulated in all clusters combined in *Arl13b^cKO^* (Fig. 3k).

Similarly, pathway analysis identified “Cilia-cShh” pathways, such as upregulated Amyloid-beta precursor protein metabolic process (Clusters 0, 1, 2 in *Arl13b^cKO^*, Clusters 0, 2 in *Ift88^cKO^*) and PI3K signaling (Cluster 0 in *Arl13b^cKO^*, Clusters 0, 2 in *Ift88^cKO^*), and downregulated ATP production (Cluster 0, 2 in *Arl13b^cKO^*, Clusters 0, 1, 2 in *Ift88^cKO^*), negative regulation of IL-1/1*β* production (Cluster 0, 2 in *Arl13b^cKO^*, Clusters 0, 1, 2 in *Ift88^cKO^*), and extracellular transport (Cluster 2 in *Arl13b^cKO^*, Clusters 0, 2 in *Ift88^cKO^*) (Fig. 3l, Extended Data Fig. 5n). Notably, in Cluster 1, the “synaptic regulation” specialized cluster, “Cilia-cShh” pathways include downregulated synaptic maturation and assembly, and upregulated negative regulation of phagocytosis, suggesting that the cShh signaling defects in *Arl13b^cKO^* Cluster 1 astrocytes result in an attenuated gene signature in synaptic regulation (Fig. 3m). Many of these “Cilia-cShh” pathways similarly altered in *Ift88^cKO^*, such as the downregulated ATP production, synapse assembly, extracellular transport, and upregulated amyloid precursor protein catabolic process, and PI3K signaling (Extended Data Fig. 5n).

In the cerebellum, the *Arl13b^cKO^* enriched BGIII cluster was found to be slightly increased in *Smo^cKO^* but not present in *SmoM2-Arl13b^cKO^* (Fig. 3n, o). BGIII marker gene expression shows a “Cilia-cShh” pattern: upregulated in both *Arl13b^cKO^* and *Smo^cKO^* and rescued in *SmoM2-Arl13b^cKO^* (Extended Data Fig. 5o, p). Thus, the enrichment of BGIII in *Arl13b^cKO^* is likely induced by cShh signaling defects. “Cilia-cShh” pathway analysis identified pathways common to BGI/II and VA clusters (e.g., metal ion transport, water homeostasis, and interleukin 1/1 *β* production) (Extended Data Fig. 5q), as well as pathways differentially altered in each cluster (Fig. 3p). Notably, unique to BGII, the cerebellar synaptic regulation associated cluster, “Cilia-cShh” pathways include upregulated regulation of postsynaptic neurotransmitter receptor activity and negative regulation of phagocytosis (Fig. 3p), suggesting that Cilia-cShh defects induced attenuated synapse regulation gene signature in BGs, similar as in cortical astrocyte Cluster 1. Together, these results revealed the differentially altered gene signatures in distinct astrocyte subtypes specifically caused by defective Cilia-cShh signaling.

### Primary cilia modulate astrocyte morphological complexity in a subtype-dependent manner independent of cShh signaling

Excluding the Cilia-cShh gene signatures, the remaining large portion of the alterations in gene expression prompted us to further define the impact of astrocyte ciliary dysfunction independent of cShh signaling. To do this, we assessed DEGs and pathways that distinguished *Arl13b^cKO^* from *Smo^cKO^* cortical astrocytes. This highlighted the downregulation of genes associated with actin dynamics regulation in *Arl13b^cKO^* but not in *Smo^cKO^* compared to controls (Fig. 4a). Similarly, we found downregulation of actin dynamics related pathways in *Ift88^cKO^* (Extended Data Fig. 6a) The GTPase RhoA is a main regulator of actin cytoskeleton that induces F-actin formation and reduced actin dynamics^79^. Indeed, we found that RhoA activity was significantly elevated in cultured *Arl13b^cKO^* but not in *Smo^cKO^* cortical astrocytes (Fig. 4b). Consistent with the elevated RhoA activity, F/G-actin ratio (Fig. 4c, d) and F-actin formation was increased in *Arl13b^cKO^* but not in *Smo^cKO^* cortical astrocytes (Fig. 4e). These results suggest that ciliary defects in astrocytes result in reduced actin dynamics in cortical astrocytes by mechanisms independent of cShh signaling.

**Fig. 4:**
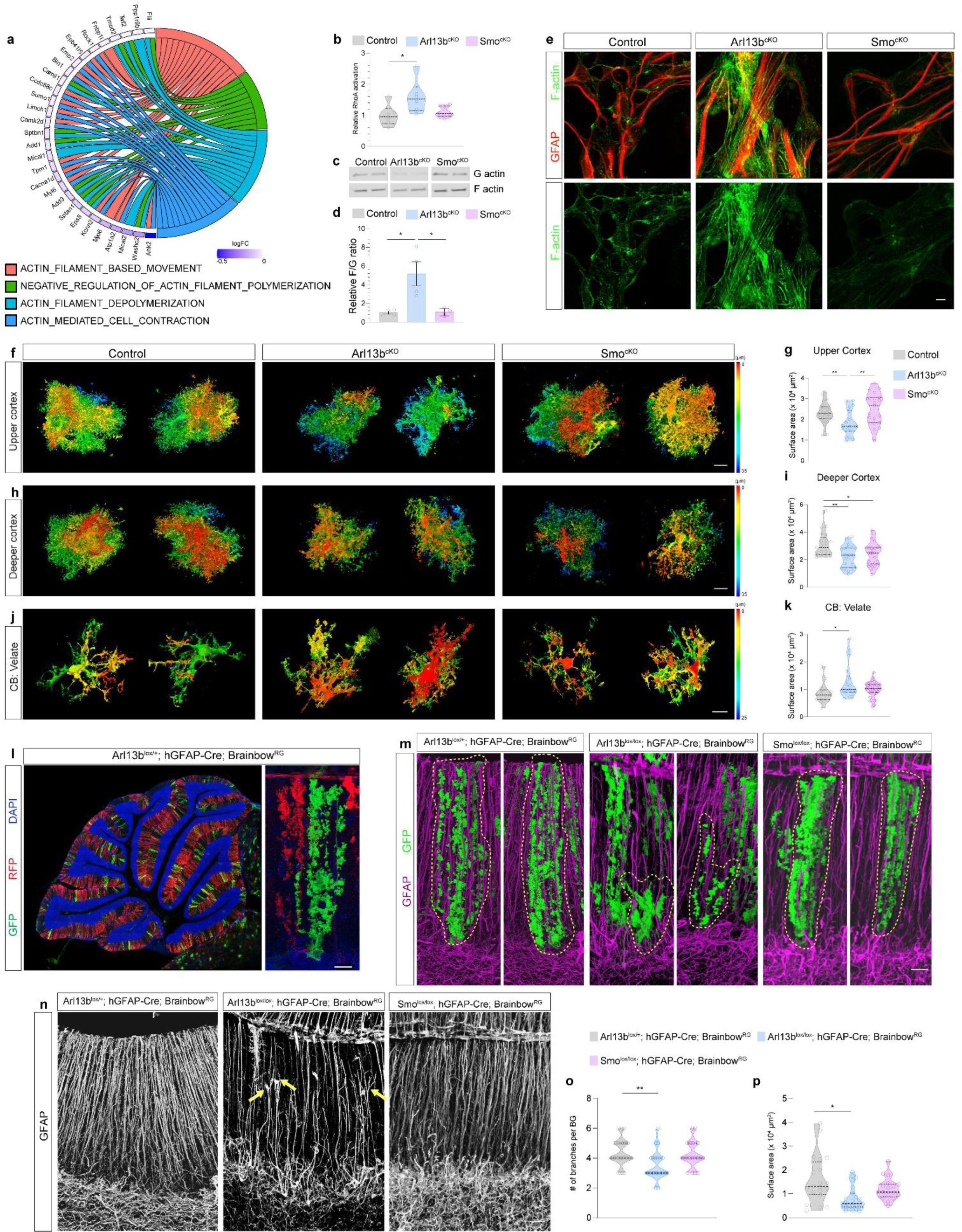
Ciliary deficient astrocytes show reduced actin dynamics and morphological changes. **(a)** Circos plot of top GO biological processes enriched for actin dynamics regulation in *Arl13b^cKO^* cortical Cluster 1. **(b)** RhoA activity is elevated in *Arl13b^cKO^* but not in *Smo^cKO^* cultured astrocytes. N= 6 per group. **(c, d)** F/G actin ratio is elevated in cultured *Arl13b^cKO^* but not in Smo*^cKO^* cortical astrocytes. **(d)** Quantification of F/G actin ratio. N= 4 biological replicates per group. **(e)** F-actin stained by phalloidin in cultured control, *Arl13b^cKO^*, and *Smo^cKO^* cortical astrocytes. **(f-k)** Representative morphology images from upper layer cortex (CTX: upper layer, f), deeper cortex (CTX: deeper layer, h), cerebellar velate astrocytes (CB: VA, j) are shown with pseudocolored depth coding. **(g, i, k)** Surface area was measured after morphology reconstruction using IMARIS. N= 18-35 astrocytes from 3 mice per group. **(l-p)** Morphology of cerebellar Bergmann glia was analyzed in *Arl13b^lox/+^; hGFAP-Cre; Brainbow^RG^* (control), *Arl13b^lox/lox^; hGFAP-Cre; Brainbow^RG^* and *Smo^lox/lox^; hGFAP-Cre; Brainbow^RG^* mice at P60. **(l)** GFP and RFP expression labels Bergmann glia in the cerebellum. **(m, n)** Bergmann glia morphology labeled with GFP (m) and GFAP (n) showed shorter Bergmann glia branches in *Arl13b^lox/lox^; hGFAP-Cre; Brainbow^RG^* brains. Yellow arrows point to the short branches failed to expand the whole molecular layer. **(o, p)** Quantifications of branch number (o) and surface area (p) of BGs are shown. N=17-26 Bergmann glia from 3 mice per group. Comparisons of three groups were performed using one-way ANOVA followed by Tukey’s test. \**P* < 0.05*, **P* < 0.01. Scale bars, 10 µm.

The actin network is a key regulator of cell morphology. Given that the morphological complexity is a cardinal feature of astrocytes critical for their function we next assessed whether ciliary mutant astrocytes show morphological defects. The AAV5-GFAP-mCherry virus was injected into the lateral ventricle in control, *Arl13b^cKO^*, *Ift88^cKO^*, *Smo^cKO^*, and *SmoM2-Arl13b^cKO^* at P0 for widespread and sparse astrocyte labeling. To assess whether Arl13b modulate astrocyte morphological regulation through cilia *vs.* extra-ciliary function, we generated AAV viral vectors that express wildtype Arl13b (AAV5-CAG-DIO-mArl13b^WT^-mCherry) and the non-ciliary mutant Arl13b^V358A^ (AAV5-CAG-DIO-mArl13b^V358A^-mCherry, localizes to the cytoplasm but maintains the GTPase activity^68, 71, 80–83^), respectively (Extended Data Fig. 6b). AAV5-CAG-DIO-mArl13b^WT^-mCherry or AAV5-CAG-DIO-mArl13b^V358A^-mCherry were injected to the lateral ventricle in *Arl13b^cKO^* at P1. By 5 weeks of age, astrocytes reach maturity and develop complex and diverse morphology with region- and subtype-specific features to meet the unique functional demands within different neural circuits^15^. In the cerebral cortex, adult protoplasmic astrocytes are highly ramified with a spongiform shape. We found that *Arl13b^cKO^* and *Ift88^cKO^* astrocytes in both upper and deeper layers show less complexity and reduced surface area (Fig. 4f-I, Extended Data Fig. 6c). By contrast, only deeper layer astrocytes in *Smo^cKO^* exhibited reduced surfaced area (Fig. 4h, i), consistent with previous studies^19, 20^. The SmoM2 expression failed to rescue the morphological defects in *Arl13b^cKO^*, consistent with our scRNAseq analysis that primary cilia modulate astrocyte morphology by cShh-independent mechanisms (Extended Data Fig. 6c). The expression of Arl13b^WT^, but not Arl13b^V358A^, led to a rescue of the morphological defects in *Arl13b^cKO^* astrocytes, suggesting that Arl13b functions in primary cilia to modulate astrocyte morphology (Extended Data Fig. 6c).

In the cerebellum, Bergmann glia (BG) and velate astrocytes (VA) show drastically distinct morphology: VAs localized in the granule cell layer show a star-shape, whereas BGs localized adjacent to the Purkinje cells are elongated and bushy-looking, extending vertical branches spanning the entire molecular layer. In contrast to the cortical protoplasmic astrocytes, in the cerebellum, *Arl13b^cKO^* VAs showed more complex morphology with increased surface area, whereas *Smo^cKO^* showed no significant changes compared to controls (Fig. 4j, k). The AAV labeling strategy was not efficient for BG labeling. To determine how primary cilia influence BG morphological development, we used *Arl13b-hGFAPCre^cKO^* and *Smo-hGFAPCre^cKO^* mice. A Cre reporter allele *Brainbow^RG 84^* that drives sparse expression of membrane-tagged fluorescent proteins (mRFP, mGFP, and mBFP) in astroglial progenitors under the *GLAST* promoter was incorporated for morphological analysis (Fig. 4l). We found that the *Arl13b-hGFAPCre^cKO^* BGs have reduced surface area, form fewer main branches that are shorter and thinner, and fail to extend the entire molecular layer (Fig. 4m-p). By contrast, *Smo-hGFAPCre^cKO^* BGs showed no significant changes (Fig. 4m-p). We analyzed both sex groups in all genotypes but did not find any sex differences in morphology (data are combined from male and female samples).

Together, these results suggest that dysfunction of primary cilia results in defects in astrocyte actin dynamics and morphological development in a subtype-dependent manner by cShh independent mechanisms. The differences between the *Arl13b^cKO^* and *Smo^cKO^* based on RhoA activity and the consequent increased levels of F-actin likely causes a more severe astrocytic morphological defect in *Arl13b^cKO^ vs. Smo^cKO^*.

### Primary cilia deficiency results in altered cell signaling states in astrocytes via cShh independent mechanisms

Among the altered gene signatures in *Arl13b^cKO^* independent of cShh signaling, elevated GPCR signaling and ER stress response pathways were common to all 3 cortical clusters (Extended Data Fig. 7a, e). *Ift88^cKO^* also shows similar changes (elevated GPCR in Cluster 0, and ER stress in Cluster 0, 1, 2) (Extended Data Fig. 3d-f). Chrm3 that encodes a GPCR M3 (Muscarinic Acetylcholine Receptor 3) was the most upregulated gene in the GPCR signaling pathway in *Arl13b^cKO^*. Immunostaining of M3 showed its significant upregulation in cultured primary cortical astrocytes in *Arl13b^cKO^* but not in *Smo^cKO^* compared to controls (Extended Data Fig. 7b). We also examined the activity of mitogen-activated protein kinase (MAPK), a convergent GPCR downstream effector signaling pathway^85^, and found a significant increase of phospho-MAPK level in *Arl13b^cKO^* cultured cortical astrocytes compared to controls (Extended Data Fig. 7c, d). These results support our scRNA seq results that primary ciliary defects lead to an elevated GPCR signaling activity in cortical astrocytes that is induced independent of cShh signaling defects.

To further explore the molecular mechanisms driving the divergence between *Arl13b^cKO^* and *Smo^cKO^*, we next examined single-cell regulatory network inference and clustering (SCENIC)-predicted transcription factor (TF) activation scores^86^. Indeed, *Arl13b^cKO^* and *Smo^cKO^* show distinct gene regulatory networks (GRNs) with core TFs unique to each other (Extended Data Fig. 7f-h). For cortical astrocytes, while developmental regulation networks like Klf13, JunB, Bmyc were robustly active in *Smo^cKO^*, GRNs highly activated in *Arl13b^cKO^* were implicated in ER stress (Xbp1, Atf6), inflammation/ stress response (Smad3, Nr4a1, Nfe2l1, Mybl1), and immediate-early response (Fos, Egr1). Immunostaining of Xbp1 and Atf6 showed enrichment within the nuclei of *Arl13b^cKO^* but not *Smo^cKO^* or control astrocytes (Extended Data Fig. 7i), confirming an exacerbated ER stress response in *Arl13b^cKO^* astrocytes^87^. Similarly, for cerebellar astrocytes, GRNs highly activated in *Smo^cKO^* BGI and VA clusters are associated with developmental regulation (Klf2, Klf9, Usf1, Usf2, Hmgb1, Smc3, and Elf1), whereas highly active GRNs in *Arl13b^cKO^* were related to inflammation (Cebpd, Hif1a, Foxj2, Irf3), ER stress (Xbp1), and energy metabolism (Ppargc1a) (Extended Data Fig. 7j, k). Together, these data highlight distinct cell signaling states and discrepant gene regulatory network activity between *Arl13b^cKO^* and *Smo^cKO^* that contribute to their distinct cellular phenotypes.

### Cilia-deficiency in astrocytes leads to profound neuronal developmental defects and behavioral deficits

Astrocytes form integral interactions with neurons to facilitate synapse formation and maturation, and synaptic pruning via phagocytosis to optimize synaptic connectivity^22, 88–91^. Given our findings that ciliary deficient astrocytes have attenuated gene signatures in synaptic regulation and morphological defects, we predicted that astrocytes deficient of primary ciliary function would exhibit compromised ability to actively modulate neuronal synapses. To test this idea, we first examined the phagocytotic abilities of *Arl13b^cKO^* cortical astrocytes *in vitro* and found a significant reduction in the amount of synaptosomes engulfed by *Arl13b^cKO^* astrocytes compared to controls (Fig. 5a, b). Further, we found a significant decline in the amount of Vglut1^+^ synaptic puncta engulfed by *Arl13b^cKO^* and *Ift88^cKO^* astrocytes both in the upper and deeper layers of the cortex, as well as the hippocampal CA1 in *Arl13b^cKO^* compared to the controls (Fig. 5c-h, Extended Data Fig. 8a-d). Interestingly, male *Arl13b^cKO^* astrocytes exhibited a more severe decline in their synaptic pruning abilities compared to female *Arl13b^cKO^* astrocytes (Fig. 5c-h, Extended Data Fig. 8b-e). Notably, the expression of Arl13b^WT^, but not Arl13b^V358A^, led to a rescue of the pruning defects in *Arl13b^cKO^* astrocytes, suggesting that Arl13b functions in primary cilia to modulate astrocyte pruning abilities (Extended Data Fig. 8f, g). Further, the pruning abilities of *SmoM2-Arl13b^cKO^* astrocytes are similarly defective compared to *Arl13b^cKO^* (Extended Data Fig. 8f, g), suggesting that primary cilia modulate astrocytes’ pruning abilities via cShh independent mechanisms.

**Fig. 5:**
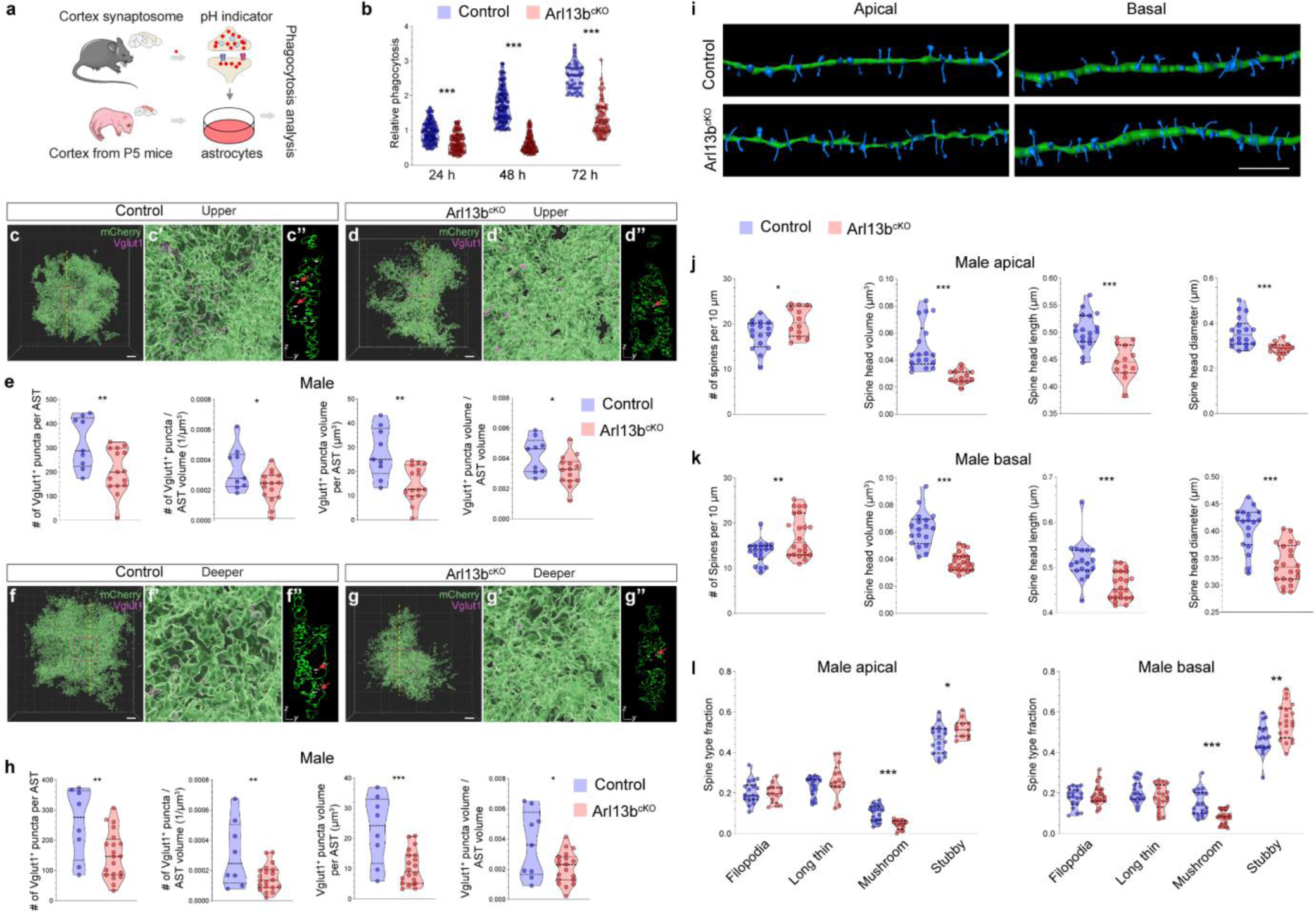
Ciliary deficient astrocytes show synaptic pruning deficits and result in neuronal spine defects. **(a)** Illustration of the *in vitro* phagocytosis assay in cortical astrocytes. **(b)** The relative phagocytosis ability of cultured control and *Arl13b^cKO^* astrocytes at 24h, 48h, and 72h post addition of synaptosomes. N= 44-135 astrocytes per group at each time point. **(c, d, f, g)** Representative images of IMARIS volumetric reconstruction of an astrocyte (mCherry, green) containing Vglut1 (magenta) in upper (c, d) and deeper (f, g) cortex in male control and *Arl13b^cKO^* brains. (c’, d’, f’, g’) Enlarged images in the red squares in c, d, f, g. (c’’, d’’, f’’, g’’) Cross-section images from the locations indicated by the yellow lines in c, d, f, g. Red arrows point to representative Vglut1 puncta engulfed in astrocytes. **(e, h)** Quantifications of the Vglut1^+^ puncta within upper- and deeper-cortical astrocytes from male control and *Arl13b^cKO^* brains. N=8-20 astrocytes from 3 mice per region per group. **(i)** Representative images of IMARIS volumetric reconstructions of apical and basal dendrites from primary motor cortex (M1) of *Arl13b^+/+^; Aldh1l1Cre^ERT2^; Thy1-eGFP* (Control) and *Arl13b^lox/lox^; Aldh1l1Cre^ERT2^; Thy1-eGFP (Arl13b^cKO^)* mice at P45. Green, dendrite; Blue, spine. (**j, k**) Quantifications of spine density, head volume, head length, and head mean diameter of M1 apical and basal dendrites from male mice. **(l)** Fraction of spine types in M1 cortex of male control and *Arl13b^cKO^* mice. N= 12-22 dendrites from 3 mice per region per group. Scale bars, 5 µm. Comparisons of two groups were performed using unpaired t-test. \**P* < 0.05, \*\**P* < 0.01, \*\*\**P* < 0.001.

To further determine whether synapse maturation is compromised by the ciliary deficiency in astrocytes, *Arl13b^cKO^* mice were bred with *Thy1-eGFP* line^92^ to label the layer V pyramidal neurons in the cortex for dendritic spine analysis. Indeed, *Arl13b^cKO^* dendrites exhibited increased spine density, but reduced spine size (Fig. 5i-l, Extended Data Fig. 8h-m). Consistent with the sex difference in synaptic pruning abilities of *Arl13b^cKO^* astrocytes, male *Arl13b^cKO^* spines (Fig. 5i-l, Extended Data Fig. 8h, i) exhibited more severe changes than females (Extended Data Fig. 8j-m). These results demonstrated that loss of cilia compromises astrocytes’ ability to modulate synapse formation and maturation, ultimately resulting in defective neuronal development.

To further assess how ciliary deficiency disrupts astrocytes’ intercellular communications in the brain and neuronal transcriptomes, we performed single nuclei RNAseq (snRNAseq) with the cortices from *Arl13b^cKO^* and control brains (Fig. 6a, Extended Data Fig. 9a-c). Using CellChat to infer alterations in intercellular signaling^93^, we analyzed astrocyte intercellular communications with excitatory neurons, inhibitory neurons, microglia, oligodendrocytes and endothelial cells. *Arl13b^cKO^* astrocytes exhibited an overall reduction in both the number and strength of outgoing signaling communications to excitatory and inhibitory neurons, and endothelial cells (Extended Data Fig. 9d-f). Remarkably, we found that 3220 genes in upper layer cortical neurons, 3115 genes in deeper layer cortical neurons, and 3454 genes in inhibitory cortical neurons were significantly suppressed in *Arl13b^cKO^*. GSEA analysis focused on the excitatory and inhibitory neurons revealed that pathways such as dendrite development, synaptic maturation and transmission, neuronal morphogenesis, Ca^2+^/ cAMP signaling and electrical coupling, were all significantly attenuated in *Arl13b^cKO^* neurons (Fig. 6b). Together, these analyses revealed a broad disturbance of astrocyte intercellular connectomes and profound changes in neuronal transcriptomic profiles induced by ciliary deficiency in astrocytes.

**Fig. 6:**
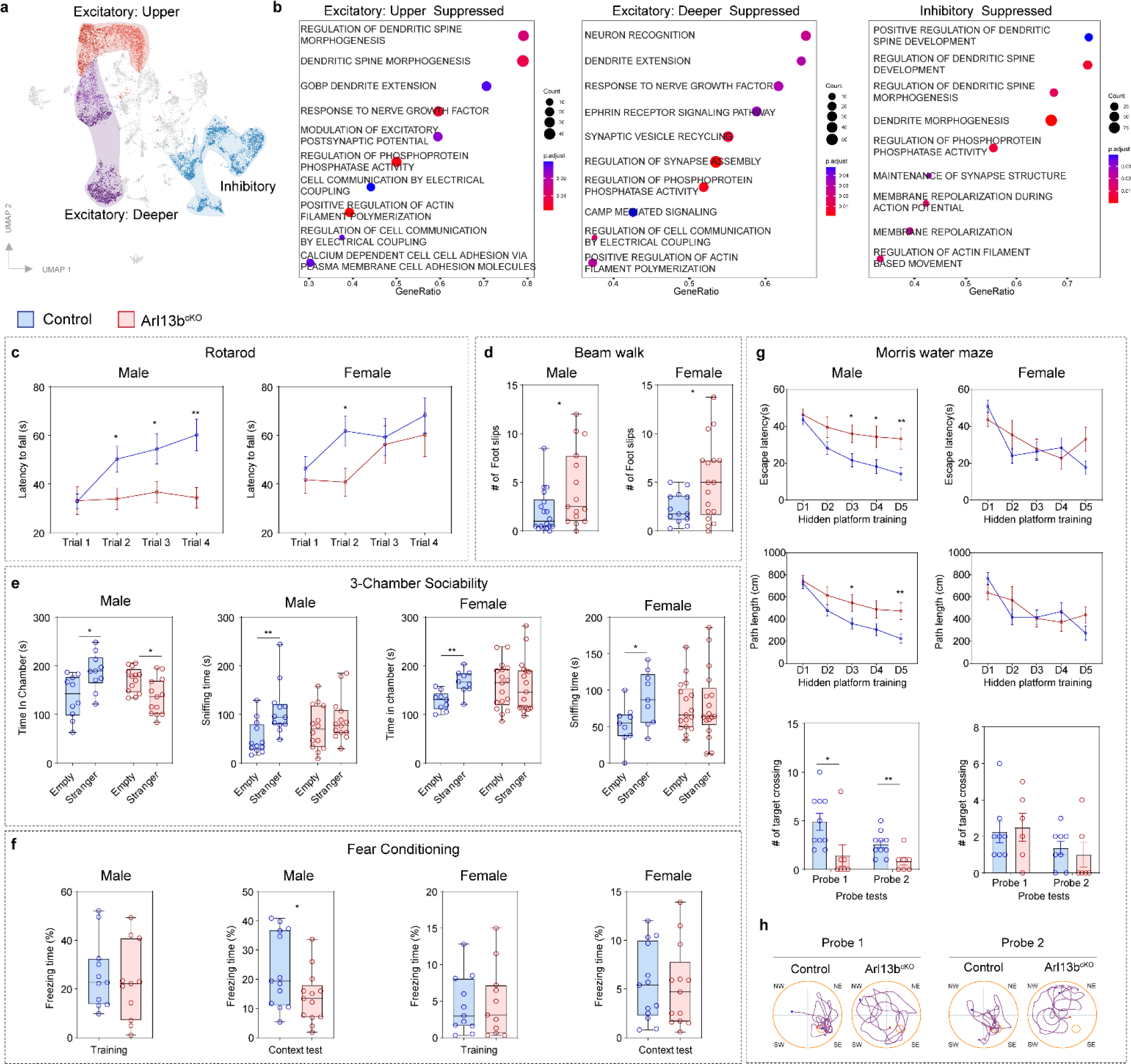
*Arl13b^cKO^* mice show disturbed neuronal transcriptomic profiles and behavioral deficits. **(a)** UMAP plot of 13670 cells of single nuclei RNAseq from the cortices of control and *Arl13b^cKO^* mice. Identified excitatory neurons (upper and deeper cortex) and inhibitory neurons are highlighted. **(b)** Top supressed GO biological processes in excitatory and inhibitory neuron in *Arl13b^cKO^* brains. **(c)** Quantifications of the latency time to fall in the rotarod test. **(d)** Quantification of the number of foot slips in the beam walk test. **(e)** Quantifications of the time in chamber and sniffing time in the 3-chamber sociability test. **(f)** Quantifications of % of freezing time in the fear conditioning test. **(g)** Quantifications for the Morris water maze test. N= 9-16 mice per group. **(h)** Representative track plots of control and *Arl13b^cKO^* male mice in the Morris water maze probe tests. Blue dot, track start; Red dot, track end. Comparisons were performed using unpaired t-test. \**P* < 0.05, \*\**P* < 0.01, \*\*\**P* < 0.001.

We next sought to assess the behavioral impact following ciliary loss in astrocytes and whether the Shh GOF leads to improved performance in *SmoM2-Arl13b^cKO^* mice. Given the sex differences in synaptic pruning and spine defects in *Arl13b^cKO^* mice, both male and female *Arl13b^cKO^*, *SmoM2-Arl13b^cKO^* and control mice were subjected to behavioral assays testing motor, sociability, learning and memory (Fig. 6c-h). Both male and female *Arl13b^cKO^* mice showed deficits in motor and sociability (Fig. 6c-e). Interestingly, only male but not female *Arl13b^cKO^* mice showed significant deficits in contextual learning, spatial learning and memory (Fig. 6f-h). Notably, *SmoM2-Arl13b^cKO^* males showed similar deficits in motor, spatial learning and memory, but significantly improved contextual learning and memory, and sociability compared to *Arl13b^cKO^* males (Extended Data Fig. 10) Thus, ciliary deficient astrocytes resulted in behavioral deficits in motor, sociability, learning and memory, with a sexual dimorphism that males showed an overall more severe deficits compared to females. Further, Shh GOF in ciliary mutant astrocytes resulted in partial functional rescue.

## Discussion

Here we show that the vast majority of astrocytes across the entire brain possess a singular primary cilium. Via canonical Shh signaling, primary cilia modulate astrocyte subtype-specific core features in synaptic regulation, intracellular transport, energy and metabolic processes. Independent canonical Shh signaling, primary cilia regulate astrocyte morphological maturation and coordinate balanced astrocyte intracellular signaling state. Consequently, primary ciliary deficient astrocytes result in neuronal developmental defects and behavioral deficits associated with human ciliopathies and related developmental disorders such as ID and ASD. Our findings illustrate a novel mechanism of how astrocytes respond to environmental modulation for diversification and maturation, by utilizing specialized signaling antennae, primary cilia. Our data also demonstrates that disrupted astrocyte development profoundly impacts brain development and function.

### The requirement of fully functional primary cilia in developing astrocytes

We used two ciliary mutants, the *Arl13b^cKO^*, in which a ciliopathy causative gene deletion leads to partially disrupted primary cilia in astrocytes, and the *Ift88^cKO^*, in which primary cilia are completely ablated in astrocytes. Consistent with the more severe ciliary disruption in *Ift88^cKO^* compared to *Arl13b^cKO^*, scRNAseq revealed significantly more differentially expressed genes (DEGs) in *Ift88^cKO^* astrocytes. However, we found striking similarities between *Arl13b^cKO^* and *Ift88^cKO^*: (i) The most enriched pathways in each cortical astrocyte clusters are similarly disturbed in *Arl13b^cKO^* and *Ift88^cKO^*; (ii) The top DEGs and pathways found in each *Arl13b^cKO^* astrocyte clusters are similarly changed in *Ift88^cKO^* clusters; (iii) *Arl13b^cKO^* and *Ift88^cKO^* astrocytes show similar morphological and pruning defects. These results suggest that although with different severity, these two ciliary gene deletions result in convergent transcriptional and cellular disturbances in developing astrocytes. Furthermore, only the wildtype Arl13b, rather than the non-ciliary Arl13b^V358A^, rescued the morphology and pruning defects in *Arl13b^cKO^*, suggesting that Arl13b functions in primary cilia to modulate astrocyte morphology and pruning abilities. To date, 5 hypomorphic mutations of *ARL13B* that lead to primary ciliary dysfunction have been identified to cause Joubert Syndrome Related Disorders (JSRD), a severe form of ciliopathies associated with ID and ASD^94–99^, suggesting that even partial loss-of-function of primary cilia can result in severe brain developmental defects and cognitive deficits. Taken together, these results demonstrate a critical and specific requirement of fully functional primary cilia in the regulation of astrocyte developmental programs.

### Primary cilia signaling modulate astrocyte subtype-specific features

Despite the importance of environmental modulation on astrocyte maturation, only a few extracellular signals have been identified, among which few has been shown to specify astrocyte diversity^11, 17, 22, 58–60^. Using scRNAseq, we analyzed major astrocyte subtypes in the cerebral cortex and the cerebellum and found that they each show distinct core features in synaptic regulation, intracellular transport, signaling states, metabolic processes, and immune reactivity. Importantly, primary ciliary defects significantly alter these features in different astrocyte subtypes, such as weakening the “synaptic regulation”, “intracellular transport”, and “ATP production” properties in cortical subtypes, and enhancing an “immune reactive” subtype in the cerebellum. These data provide a novel cellular mechanism and molecular insights into how astrocytes receive environmental cues, via their primary cilia, to specify their subtype-specific and region-specific developmental programs across the brain.

In addition to genetic features, we found that primary cilia are important regulators of astrocyte morphological maturation in a subtype-dependent manner: Ciliary mutant protoplasmic astrocytes in the cerebral cortex and the hippocampus, as well as mutant Bergmann glia in the cerebellum, show significant reduction in size and complexity. In contrast, the VA astrocytes in the cerebellum show increased size and complexity. BGs closely interact with Purkinje cells whereas VAs mainly interact with granule cells. It is possible that Purkinje cells and granule cells secrete different cues to differentially influence their astrocyte partners. This is supported by recent evidence that Purkinje cells but not granule cells secrete Shh, therefore enabling BGs but not VAs to be exposed to higher concentrations of Shh^11^. Interestingly, we found that primary cilia modulate astrocyte morphology via cShh independent mechanisms, likely through the regulation of RhoA GTPase and actin dynamics. Future studies using astrocyte subtype labeling and isolation methods are needed to explore further how local environment drives different maturational programs and functional outcomes for distinct astrocyte subtypes on the basis of differences in local cues and primary cilia signaling cascades. It also remains a question whether defective cilia can elicit cell-autonomous changes on astrocyte morphology. Future work identifying specific ciliary signaling in astrocytes, as well as how structurally defective primary cilia may alter cytoskeletal dynamics and organization would be important for more mechanistic insights underlying the primary cilia-based astrocyte developmental regulation.

### Profound neuronal defects, ASD and ID relevant behavioral deficits can be induced by ciliary deficient astrocytes

A cardinal feature of astrocytes is their extensive morphological complexity that enables their close interactions with neurons and other cell types in the brain^22, 88–91^. Astrocyte morphology defects could lead to insufficient cell-cell contact surface, signaling strength, and lack of morphological plasticity required for the optimal astrocyte-neuron communications to compromise neuronal development. Indeed, astrocyte morphological defects have been shown to be associated with circuit malfunction and common CNS disorders^66^. In addition to morphology, the phagocytosis and synaptic pruning abilities of astrocytes are critical for neuronal circuits maturation^7, 10, 100, 101^. We found that ciliary deficient astrocytes show morphology defects and reduced synaptic pruning abilities *in vivo*. Consequently, dendritic spines show maturational defects. Further, single-nuclei RNAseq analyses revealed that over 3000 genes enriched in pathways including synaptic maturation and neuronal plasticity are significantly suppressed in both excitatory and inhibitory neurons. Ultimately, mice with primary ciliary deficient astrocytes exhibit behavioral deficits associated with human ciliopathies and related developmental disorders including ASD and ID. Although dysregulated astrocytes are associated with neurodevelopmental disorders such as autism spectrum disorders (ASD) and intellectual disabilities (ID), causal evidence linking the disruption in astrocyte development with neuronal defects and behavioral consequences related to ASD and ID remains sparse^27, 28, 91, 102^. Our findings provide compelling causal evidence to demonstrate how disruption in astrocyte development can profoundly compromise neuronal programs to manifest behavioral deficits underlying neurodevelopmental disorders.

Previous studies dissecting the function of primary cilia in brain development and malformations associated with ciliopathies have been primarily focused on neuronal primary cilia and how they cell-autonomously regulate neuronal development and function^34, 36, 40–42, 51–57, 103–106^. Our finding here provides a new way to think about how primary ciliary deficiency could exert profound impact on brain development and health, by acting on glial cells such as astrocytes to modulate the mutual neuron-astrocyte communications and indirectly regulate neuronal programs. Future studies are needed to further delineate how ciliary defects in glial cells are contributing to the pathology of ciliopathies and related brain disorders.

### Primary cilia mediate Shh signaling and much beyond to impact astrocytes

Using comparative scRNAseq analyses between two ciliary mutants, Shh loss- and gain-of-function mutants, we provide novel insights: 1) The cShh-dependent modulation in astrocytes is at least in part mediated through primary cilia; 2) All cortical protoplasmic and cerebellar astrocytes, despite differences in cShh signaling activity, use cilia-cShh signaling to differentially modulate their subtype-specific core features; Thus, our results for the first time directly establish a functional link of how primary cilia mediate cShh signaling in astrocytes to modulate their developmental programs. 3) We show that cShh signaling only account for <50% of the genetic changes in ciliary mutant astrocytes. Independent of cShh signaling, regardless of subtype or region, cilia dysfunction disrupts astrocyte morphology, signaling balance, and triggers ER stress response, revealing a conserved, cShh independent ciliary function in astrocyte cytoskeletal regulation, cell signaling, and homeostasis across the brain. 4) Shh gain-of-function in ciliary mutant astrocytes resulted in rescue of the behavioural deficits in contextual learning and memory, and sociability, but not in motor functions or spatial learning and memory. Together, these results highlight that primary cilia mediate both Shh-dependent and - independent pathways to modulate astrocyte subtype-specific developmental programs.

The overall elevated cellular signaling activities in ciliary mutant astrocytes, as evidenced by scRNAseq GSEA analysis (Fig. 2, augmented “GPCR signaling activity”, “response to external stimuli”, and increased phospho-MAPK level), points to an intriguing scenario that primary cilia transduce “inhibitory signaling” to restrict and fine tune astrocytes’ signaling activities in response to external stimuli. Heightened cellular signaling in response to external stimuli can increase biosynthetic and energy demand and oxidative stress, which induces accumulation of misfolded proteins and consequently trigger the activation of ER stress and stress/inflammatory response pathways^87, 107^. This is particularly exemplified in the cerebellum, where ciliary dysfunction significantly enriched a subtype of BGs (BGIII) that shows an energy deficient and stress/immune reactive signature that is barely present in control brains. We postulate that primary ciliary defects not only disturb the communication between astrocytes and their surrounding environment for instructional information but may also lead to broad, long-term consequences that compromise cellular metabolism and homeostasis. Such ciliary signaling function might be widely applicable to other cell types and organ systems considering the nearly ubiquitous presence of primary cilia in the body and the wide variety of biological processes and disease conditions they are involved in^36, 108, 109^.

Compared to canonical Shh signaling, much less is understood about how primary cilia transduce other signaling pathways and their cellular outcomes in each cell type. To date, a few dozen signaling receptors including GPCRs and receptor tyrosine kinases have been collectively reported to localize to primary cilia of a variety of cell types, including neurons^32, 34, 41^. Indeed, our scRNAseq analysis revealed that both ciliary mutants show significantly altered GPCR and RTK signaling and related pathways (e.g., MAPK, PI3K, TGF-β) in distinct cortical and cerebellar astrocyte subtypes. However, little knowledge exists about the primary ciliary protein composition or cilia-signaling in astrocytes. Future work is needed to pinpoint the upstream non-Shh signaling pathways emanating from primary cilia in astrocytes. With the recent advancements in proximity labeling-based proteomics and ciliary purification techniques^110–112^, we anticipate that successful application of such methods on the comprehensive profiling of astrocytic cilia will greatly aid further investigations on how primary cilia sense and convey environmental signals to modulate astrocyte biology, and how this process is perturbed to manifest pathogenic changes in ciliopathies and associated diseases such ID, ASD, and epilepsy. In addition to neurodevelopmental disorders such as ASD and ID, astrocyte dysfunction and ciliary defects have been independently associated with an overlapping spectrum of neurodegenerative diseases such as Alzheimer’s disease and Parkinson’s disease^3, 23, 36, 113–119^. Intriguingly, elevated ER stress response that is found in ciliary mutant astrocytes, has been strongly linked to Alzheimer’s disease pathology^120^. It would be thus interesting to explore whether primary cilia defects in adult or aging astrocytes contribute to the progression of neurodegenerative disorders such as Alzheimer’s disease.

Further, it would be also interesting to examine whether primary cilia are required in mature astrocytes to detect the injury/disease triggered cues to help configure astrocyte reactions under pathological conditions. This work opens up future avenues for investigations into the cilia-mediated extracellular signals that configure astrocyte heterogeneity, and the potentially broad role of primary cilia in astrocyte signaling, metabolism, neuron-astrocyte communications, and inflammatory response underlying brain development, homeostasis and disease progression.

## Methods

### Animals

*Mice* were cared for according to animal protocols approved by the University of Calgary. The genetic mouse models used in this study were generated based on the following lines: *Arl13b^lox/lox^* (*Arl13b^tm1Tc^* /J, JAX Stock: #031077)^61^*, Ift88^lox/lox^* (*Ift88^tm1Bky/J^*, JAX Stock: #022409) ^62, 63^, *Smo^Lox/Lox^* (*Smo^tm2Amc^*/J, JAX Stock: #004526)^121^*, Aldh1l1-Cre^ERT2^* [*B6N.FVB-Tg(Aldh1l1-cre/ERT2)1Khakh*/J, JAX Stock: # 031008]*, hGFAP-Cre* [*B6.Cg-Tg(GFAP-cre/ERT2)505Fmv*/J, JAX Stock: 012849]^122^, *Ai9* [*B6.Cg-Gt(ROSA)26^Sortm9(CAG–tdTomato)Hze^*/J, JAX Stock: # 007909]^123^, *Brainbow^RG^*^124^, *SmoM2* [*Gt(ROSA)26Sor^tm1(Smo/EYFP)Amc^*/J, JAX Stock: # 005130]^125^. The following crossed in-house mouse strains were used: *Arl13b^lox(+)/lox(+)^*; *Aldh1l1-Cre^ERT2^*; (*Ai9), Ift88^lox(+)/lox(+)^*; *Aldh1l1-Cre^ERT2^*; (*Ai9), Smo^lox(+)/lox(+)^*; *Aldh1l1-Cre^ERT2^; (Ai9), SmoM2^lox/+^;* Arl13b^lox^*^(+)/lox(+)^*; *Aldh1l1-Cre^ERT2^; (Ai9)*, *Arl13b^lox(+)/lox^*; *hGFAP-Cre*; (*Brainbow^RG^), Smo^lox/lox^*; *hGFAP-Cre*; *(Brainbow^RG^)*.

### Primary astrocyte culture, transfection, and Smo localization analysis

*Arl13b^lox/lox^; Aldh1l1-Cre^ERT2^* or Arl13b^+/+^; Aldh1l1-Cre^ERT2^ mice were injected with Tamoxifen (20mg/mL, 10uL/pup) from P2 to P4. Dorsal cortices were dissected Hank’s buffered salt solution (HBSS) at P5 for astrocyte culture. The culture protocol was modified based on the published method^126^. In brief, dorsal cortices were cut into <1mm^3^ pieces and dissociated in the enzyme buffer (1 x EBSS media, 0.36% D(+) glucose, 26mM NaHCO_3_, 0.5mM EDTA) containing Papain (∼34 units/mL, Worthington), DNase I (100 units/ml, Sigma), N-Acetyl-L-cysteine (0.32 mg/mL, Sigma) and HEPES (10mM) for 1 hour in the 37 °C cell culture incubator (5% CO_2_, 95% air). Tissue pieces were gently triturated after washing using protease inhibitor stock solution [1x EBSS, 0.36% D(+)-glucose, 26mM NaHCO_3_, 4mg/mL Trypsin inhibitor, and 4mg/mL BSA, pH= 7.4]. Then, the cells were spun down and plated on poly-D-lysine-coated plastic coverslips in a Neurobasal-DMEM-based serum free medium containing 50% DMEM (Gibco), 50% Neurobasal medium, Pen/Strip (100 units/mL, Gibco), Sodium Pyruvate (1mM, Sigma), L-Glutamine (Sigma, 2 mM), HBEGF (5ng/mL), N-Acetyl-L-cysteine (5 µg/mL, Sigma), Transferrin (100 µg/mL), BSA (100 µg/mL), Putrescine (16 µg/mL), Progesterone (4nL/mL) and Sodium Selenite (0.1 µL/mL). We replaced half of the volume with fresh medium every 4 days to maintain the cultures.

Cultured astrocytes were transfected with Smo::eGFP^127^ plasmids at 12 days post culture using jetOPTIMUS® DNA Transfection Reagent (Polyplus) according to the manufacturer’s protocol. Cells were treated with SHH (100 ng/mL) on next day post transfection and fixed after another 24 hours for immunostaining and imaging analysis.

### *In vitro* astrocyte synaptic phagocytosis assay

Synaptosomes purification and pH-indicator conjugation were performed according to the published method^128^. Briefly, cortices from two mice at the age of 2 months were dissected and homogenized. Synaptosome was isolated by centrifuging the homogenate using discontinuous sucrose density gradient media (bottom to top: 23%, 10%, 3%). The pH indicator was conjugated to synaptosome in 0.1M Na_2_CO_3_ for 2 hours at room temperature in a twist shaker with gentle agitation at 30-40 rpm. The pH-conjugated synaptosomes were stored at the concentration of 2 mg/mL in −80 °C.

The *Aldh1l1Cre^ERT2^* (control) and *Arl13b^lox/lox^; Aldh1l1Cre^ERT2^* (*Arl13b^cKO^*) mice injected with tamoxifen from P2 to P4 were used to culture cortical astrocytes at P5. Astrocytes were cultured in 35-mm glass bottom dishes coated with poly-D-lysine for 10 days. The medium in the dishes of cultured astrocytes were collected in falcon tubes for use as conditional culture medium. Astrocytes were washed with DPBS three time, then treated with 10 uL pH indicator conjugated synaptosome suspended in the 400 µL conditional medium and incubated in 37 °C cell culture incubator for 40 min. The media with unbound pH indicator-conjugated synaptosomes was removed and astrocytes were washed three times with DPBS. Culture medium (1:1 of conditional and fresh medium) were added to the astrocytes for additional incubation. The engulfment of the synaptosomes by astrocytes were analyzed by live bright field and fluorescent microscopy at 24h, 48h, and 72h post synaptosome treatment. The intensity of pH indicator within each cell was calculated to reflect the phagocytosis ability of astrocytes.

### Astrocytic morphology and pruning analysis

The *Aldh1l1Cre^ERT2^* (control), *Arl13b^lox/lox^; Aldh1l1Cre^ERT2^* (*Arl13b^cKO^*), and *Smo^lox/lox^; Aldh1l1Cre^ERT2^* (*Smo^cKO^*) mice were injected with AAV5-Gfap-mCherry in the lateral ventricle at P0, and tamoxifen (20mg/mL, 10uL/mouse) were administrated from P7 to P9. For morphology analysis, mice (P35) were perfused with PBS and 4% PFA, and brains were collected and sectioned at 50 µm. Sections were stained with anti-RFP antibodies (1:1000, ROCKLAND) to label the astrocytes expressing mCherry. Confocal images were taken using a Zeiss LSM 880 Airyscan Confocal Laser Scanning Microscope and processed by ZEN black system. The quantification of surface area of astrocytes was measured by IMARIS software.

For astrocytic pruning analysis, brains were collected at P35 and cryo-sectioned at 20 µm after dehydration using 30% sucrose. Sections were stained with Vglut1 and RFP antibodies and secondary antibodies. Z-stacks confocal images were taken at 63x objectives using a Zeiss LSM 880 Airyscan Confocal Laser Scanning Microscope and processed by ZEN black system. The images were imported in IMARIS software for the engulfment analysis^129^. First, volumetric reconstructions of the astrocytes labeled with mCherry were created using the ‘Surface’ function in IMARIS after running the ‘enhance contrast’ with a mean filter of pixel radius at 1.5 µm. The following surface object reconstruction steps were performed following the guided creation wizard. The discontinuous parts of the surface not clearly traced were removed according to the fluorescence. Next, the astrocyte surface was used as the region of interest to create a mask of Vglut1 channel, defined by the signal included within the surface. New surface objects of Vglut1 that completely internalized in astrocytes were generated using the masks. Finally, the volume of the astrocytes, the volume and number of internalized Vglut1 puncta were collected from the statistics tab in IMARIS. The number/volume of Vglut1 puncta per astrocyte, and the normalized number/volume of Vglut1 puncta to the astrocyte volume were calculated.

### Immunostaining for brain sections and cultured cells

Brain sections and cultured cells were fixed by 4% PFA were immunolabeled as previously described^34, 36–42^ with the following primary antibodies: anti-RFP (rabbit, 1:500; Rockland), anti-GFP (chicken, 1:2000), anti-Arl13b (mouse, 1:500; Abcam), anti-GFAP (chicken, 1:1000; Abcam), anti-Aqp4 (rabbit, 1:500; Alomone Labs), anti-Kir4.1 (rabbit, 1:500; Millipore Sigma), anti-Xbp1 (rabbit, 1:500; Novus Biologicals), anti-ATF6 (mouse, 1:1000; Novus Biologicals), Vglut1 (Guinea pig, 1:1000; Synaptic Systems GmbH). AlexaFluor 488, 568 or 647-conjugated (Thermo Fisher) secondary antibodies were used. Nuclei were counterstained with DAPI (Sigma). For F-actin staining in cultured astrocytes, AlexaFluor 647 conjugated Phalloidin (Thermo Fisher) was used along with the secondary antibody incubation.

### G- and F-actin analysis, and RhoA activity analysis

Cortical astrocytes were cultured from the P5 mice and were collected at day 14 post culture. RhoA activity was detected using the RhoA G-LISA Activation Assay Kit (Cytoskeleton) according to the manufacturer’s protocol. G- and F-actin was detected by G-Actin/F-Actin In Vivo Assay Biochem Kit (Cytoskeleton) according to the manufacturer’s protocol. The G- and F-actin was detected by western blotting and the intensity of bands was measured by ImageJ to calculate the F/G ratio.

### Astrocyte isolation for single cell RNA seq

*Aldh1l1-Cre^ERT2^* (Control), *Arl13b^lox/lox^; Aldh1l1-Cre^ERT2^* (*Arl13b^cKO^*), *Ift88^lox/lox^; Aldh1l1-Cre^ERT2^* (*Ift88^cKO^*), *Smo^lox/lox^; Aldh1l1-Cre^ERT2^* (*Smo^cKO^*), and *SmoM2-eYFP; Arl13b^lox/lox^; Aldh1l1-Cre^ERT2^* (*SmoM2-Arl13b^cKO^*) mice were administrated with tamoxifen (20mg/mL) from P7 to P9. Cortices and the cerebellum were dissected at P52∼55. Samples from two male mice of each genotype were combined. Tissue was cut into small pieces and digested in 5mL enzyme buffer (1 x EBSS media, 0.36% D (+) glucose, 26mM NaHCO_3_, 0.5mM EDTA) containing papain (∼34 units/mL, Worthington), DNase I (20 units/ml, Sigma) for 30 min at 37 °C in a cell culture incubator (5% CO_2_, 95% air). Tissue pieces were then gently triturated into single cells after stopping the enzyme reaction by adding 10mL DMEM (Gibco) supplied with 10% FBS (Gibco). Cells were spun down and resuspended in 13mL of Percoll solution [23% Percoll (Sigma), 36mM NaCl in PBS], and carefully topped with 8mL of PBS. Cells were then centrifuged at 900g (accelerate: 4, break: 0), resuspended and washed once with FACS buffer (PBS supplied with 3% FBS). Blocked with FcBlock (Biorad) for 5 min at room temperature, cells were then stained with ACSA2-PE (Miltenyi Biotec) for 30min at 4 °C, followed by Viability Dye eFluor™ 780 (eBioscience) incubation for 5 min at room temperature. Cells were then washed with PBS solution supplied with 3% FBS (Gbico) once, and live ACSA2^+^ cells were sorted using flow cytometry. The RNase inhibitor (Sigma) was supplied in the solutions from the staining to the end of sorting.

### Single cell-RNA seq library construction, sequencing and alignment

A total of 15, 000 single cells of each sample were loaded for partitioning using 10x Genomics NextGEM Gel Bead emulsions (3’ gene expression kit, version 3.1) and processed according to the manufacturer’s protocol. The PCR amplification steps were run 12x. TapeStation D1000 was used to check the quality control (QC) of the resulted libraries. Illumina Sequencing was performed on NovaSeq SP and Illumina NovaSeq S2 at the Centre for Health Informatics (CHGI) at University of Calgary to achieve a sequencing depth of over 25,000 read pairs per cell post-aggregation. Alignment to the reference genome was performed using the CellRanger 3.1.0 pipeline^130^. Aggregated cortical samples recovered 22307 cells (Control: 4511 cells, *Arl13b^cKO^*: 3366 cells, *Ift88^cKO^:* 5099 cells*, Smo^cKO^*: 5162 cells, *SmoM2-Arl13b^cKO^*: 4169 cells), and aggregated cerebellar samples recovered 6541 cells (Control: 2193 cells, *Arl13b^cKO^*: 1595 cells, *Smo^cKO^*: 2052 cells, *SmoM2-Arl13b^cKO^*: 701 cells).

### Single nucleus RNA-seq library construction, sequencing and alignment

Nuclei were isolated from mouse (P52) cortices with cell fixation and nuclei fixation kit (Parse Biosciences). Cortices of two male mice were combined in each group. A total of 10,000 cells per group were used for library construction using the Evercode^TM^ Whole Transcriptome Mini kit (Parse Bioscience) as per the manufacture’s protocol. TapeStation D1000 was used to check the quality of the libraries. Alignment was performed with the pipeline provided by Parse Bioscience. 8914 cells and 7257 cells were recovered from *Arl13b^cKO^* (Arl13b^lox/lox^; Aldh1l1-Cre^ERT2^, tamoxifen: P7 ∼ P9) and control *(Arl13b^+/+^; Aldh1l1-Cre^ERT2^*, tamoxifen: P7 ∼ P9), respectively.

### Single cell/nucleus RNA-seq computational analysis and workflows

HDF5 matrices from aggregated datasets of single cell RNA-seq, or the processed gene expression matrices of single nucleus RNA-seq were imported into the R package Seurat (version 4.3.0) for analyses. Cells with more than 25% of mitochondrial reads were removed during the quality control step. Gene expression was normalized using sctransform^131^ R package and batch effects were corrected using Harmony^132^. Cell types were annotated manually according to the expression of cell type-specific marker genes. Cell markers and differentially expressed genes (DEGs) were identified using the FindMarker or FindAllMarker functions in Seurat. Sample-specific perturbation scores were calculated by summing DEG FCs for each astrocyte cluster^133^ and shown using Nebulosa^134^ (version 1.8.0) generated density plots. Gene set enrichment analysis to identify GO terms enriched in each cell cluster or sample was performed using GSVA R package. SCENIC^135^ was employed to identify the group-specific regulons in astrocyte clusters. CellChat^136^ was used to identify the cell-cell interaction networks for the single nucleus RNA-seq dataset.

### Mouse behavioral test

To assess the motor balance and coordination of mice, rotarod and beam walk tests were performed. For rotarod test, latency time to fall from an accelerating rod (4 ∼ 40 rpm, up to 5 min) was measured for each mouse. A total of four trails were performed for each mouse in one day. For beam walk, the video of mice crossing a 1-meter beam was taken. Four tests of each mouse were performed. The time to cross the beam and the number of foot slips were measured, and the average time and foot slips of four tests were calculated.

For fear conditioning assays, mice were tested in a conditional fear task over two days. In brief, mice received 3 pairings consisting of a light and a co-terminating 0.5mA shock of 1 second duration at day 1. Mice were tested for contextual fear memory of light. The freezing time was measured.

Sociability of mice were assessed using 3-chamber social approach. Mice first received 10min habituation to the two empty cups, and sociability was assessed for 10 min with a cup containing a sex- and age-matched stranger mouse and an empty cup. Time in chambers and the sniffing time were measured.

Morris water maze (MWM) test^137^ was used to assess spatial learning and memory of mice. In brief, mice received visible training at day 1 and invisible training from day 2 to day 6. Probe tests were performed at day 7 and day 14 to assess the short-term and long-term memory.

### Statistical analysis

All statistical tests except for single cell transcriptomics analysis were performed in Graphpad Prism 8 or R. For comparisons of two groups, unpaired two-tailed Student’s t-test was used. One-way ANOVA tests followed by Tukey’s tests were used for comparisons of three groups. Data are shown as mean ± SEM in Student’s t-test or ANOVA tests. Individual values were shown in violin plots. *P* values was considered significant when < 0.05. \**P* < 0.05, \*\**P* < 0.02, \*\*\**P* < 0.001.

## Data and code availability

Single-cell RNA-seq data is deposited at GEO and publicly available (GSE253643). Analysis code generated in this study is available through https://github.com/TheGuoLab/AstrocytesPrimaryCilia. Any additional information required to reanalyze the data reported in this paper is available from the lead contact upon request.

## Acknowledgments

J.G. is a New York Stem Cell Foundation-Robertson Investigator. This research was supported by The New York Stem Cell Foundation (NYSCF-R-N163 to J.G), the Canadian Institutes of Health Research (RN418797-438409 to J.G.), the National Science and Engineering Research Council (RGPIN-2019-04820 to J.G.). We thank the support from Harley N. Hotchkiss-Samuel Weiss Postdoctoral Fellowship (L.W.), Cumming School of Medicine Postdoctoral Scholarship (L.W.), and ACHRI STEP Postdoctoral Fellowship (L.W.).

## Author contributions

Conceptualization, L.W. and J.G.; mouse experiments and husbandry, L.W., Q.G., X. Z., V.H., B.W., M.M., S.C., C.C., and J.G.; laboratory experiments, L.W., Q.G., X. Z., V.H., B.W., A.Y., N.R., E.L., C.C., M.M., S.C., K.G., J.H., D.E., T.R., H.K., and J.G.; bioinformatics, L.W., S.A., N.R., Q.Z.; manuscript editing and review, L.W., G.G., J.B., and J.G.; manuscript writing, L.W. and J.G.

## Competing interests

The authors declare no competing interests.

---

**Extended Data Fig. 1.**
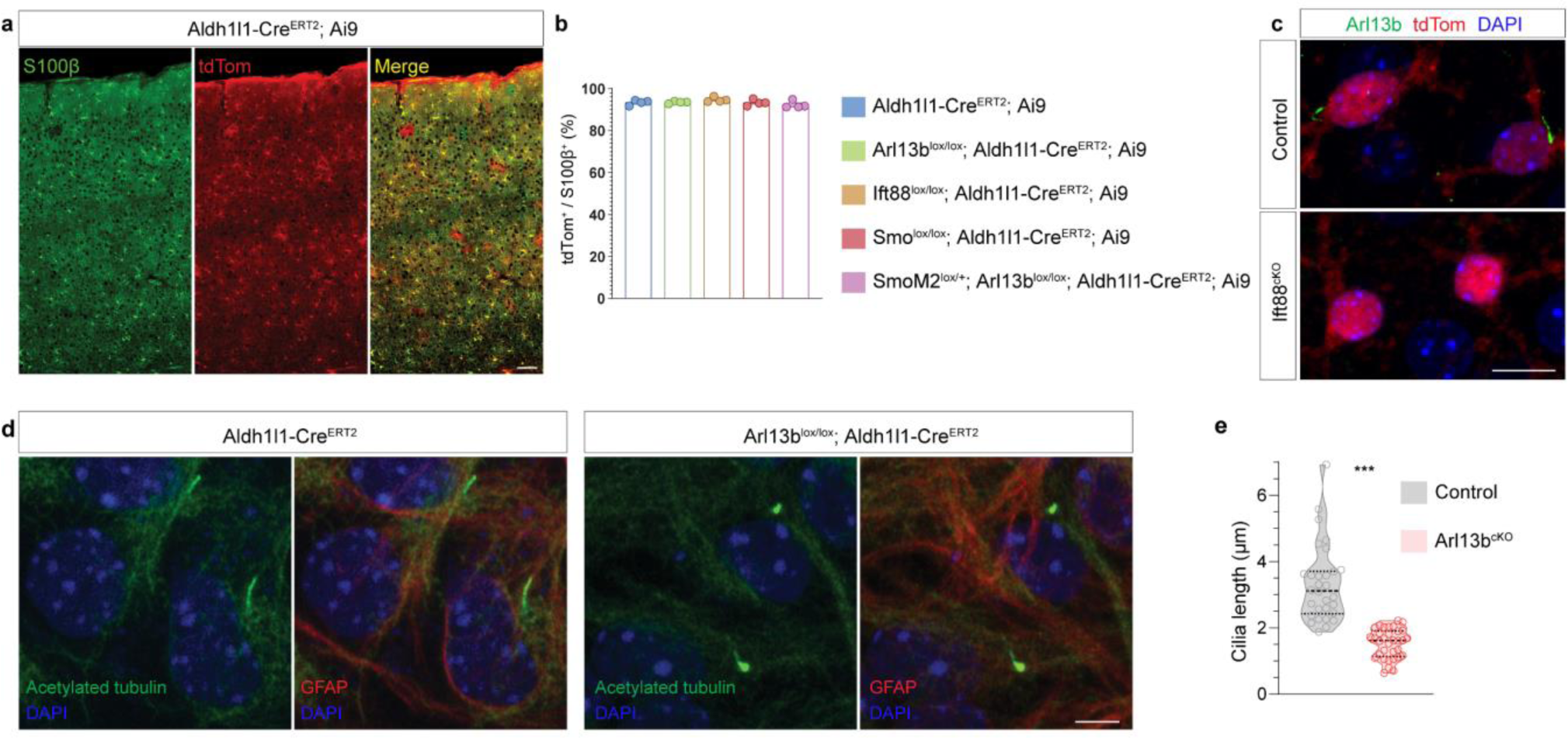
Tamoxifen mediated Cre recombination and ciliary gene deletion in astrocytes. **(a, b)** Tamoxifen mediated highly efficient and consistent Cre recombination in all 5 mouse lines. **(a)** Representative images of astrocytic tdTom (Ai9) expression in S100β^+^ astrocytes in the cerebral cortex in control brains. Mice were injected with tamoxifen from P7 to P9, and brains were collected at P35. **(b)** Quantifications of the % of S100β^+^ astrocytes that expressed tdTom in all 5 mouse lines. N=4 mice per group. **(c)** Primary cilia (labeled by anti-Arl13b) are absent in *Ift88^lox/lox^; Aldh1l1Cre^ERT2^; Ai9* (*Ift88^cKO^*) mice. Mice were injected with tamoxifen from P7 to P9, and brains were collected at P35. **(d, e)** Cilia morphology of control and *Arl13b^cKO^* cultured primary astrocytes stained by anti-acetylated tubulin antibody. N=30-33 cells per group. Comparison was performed using unpaired t-test. \*\*\**P* < 0.001. Scale bar in (a), 50 µm; scale bars, 10 µm (c), 5 µm (d).

**Extended Data Fig. 2.**
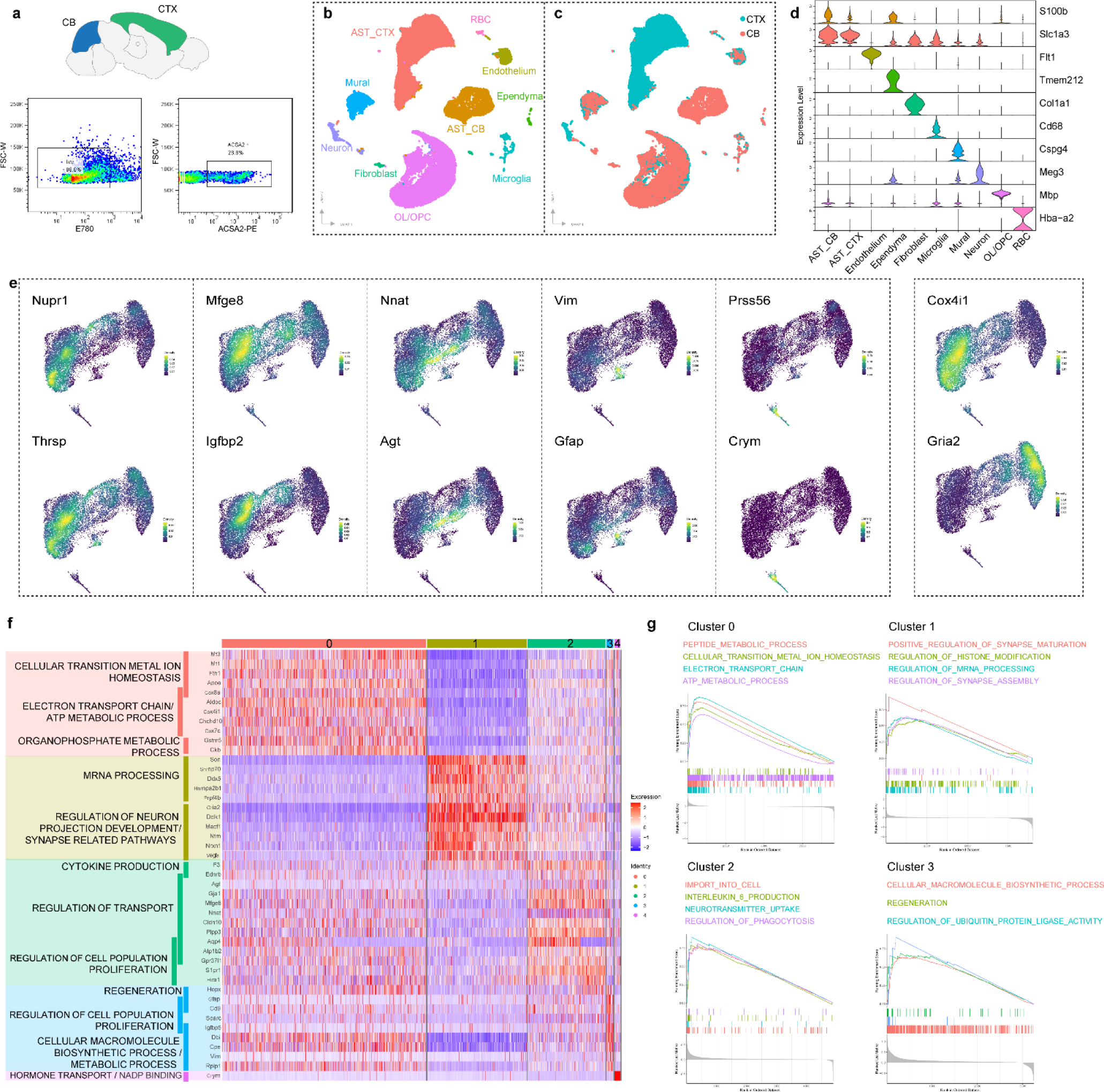
Single cell RNAseq analyses. **(a)** Astrocyte sorting using flow cytometry. Cortices and cerebellums were dissected and were digested to isolate single cells. Cells were stained with viability dye E780 and ACSA2-PE. ACSA2 positive live cells were sorted to construct the single cell RNA seq libraries. **(b)** UMAP plots of all cells annotated with main cell types. **(c)** UMAP plots of all cells grouped by tissue types from which the cells are isolated from. **(d)** Expression of representative cell type markers used for cell annotation. **(e)** Expression patterns of regional markers and representative cluster marker genes of cortical astrocytes. **(f)** Heatmap of top cortical astrocyte cluster markers and the related GO terms. **(g)** GSEA plot of enriched key GO pathways in cortical astrocyte clusters.

**Extended Data Fig. 3.**
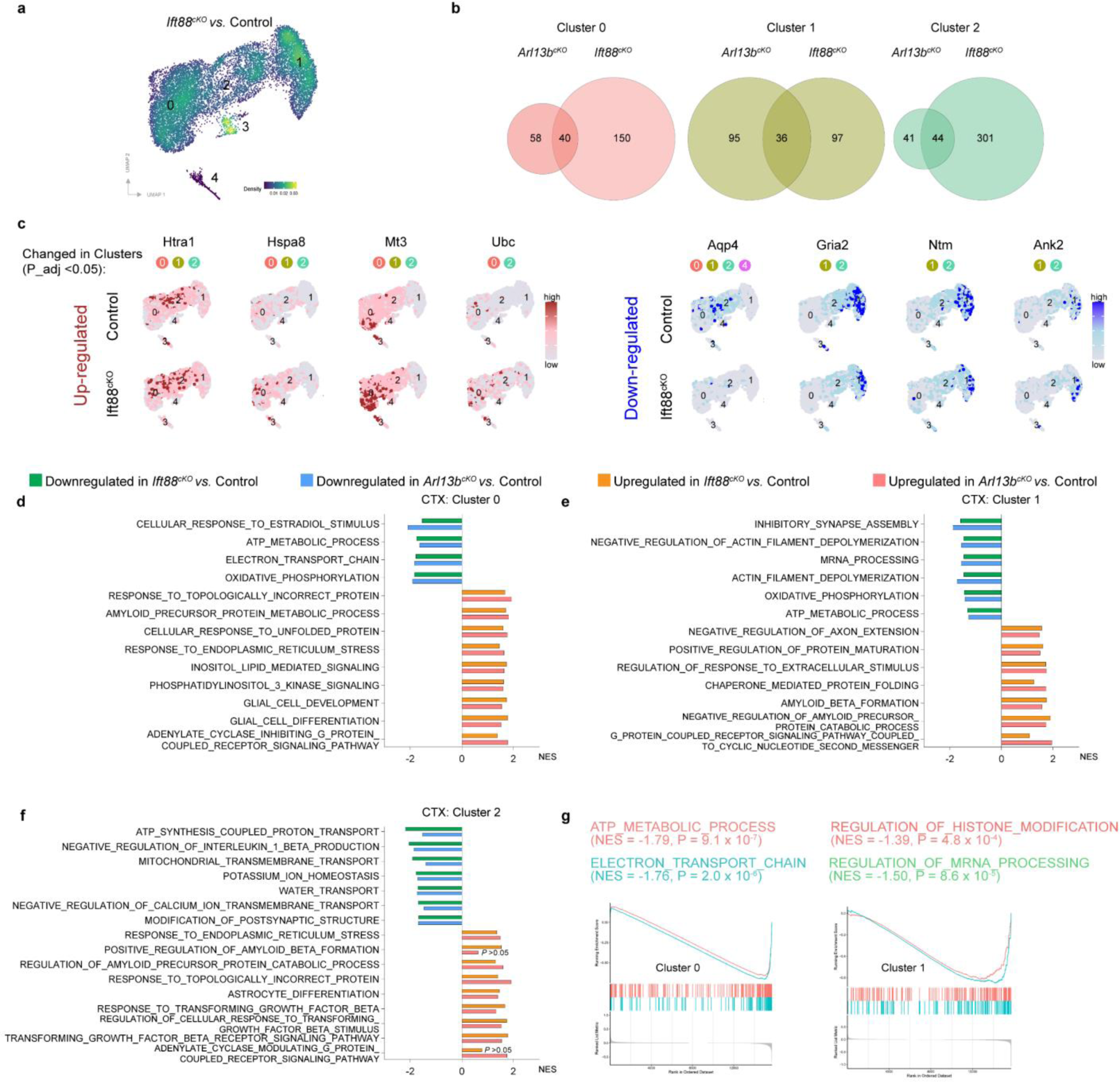
Single cell RNAseq analyses of *Ift88^cKO^* cortical astrocytes. **(a)** Kernel density estimates depicting magnitude of the molecular changes in *Ift88^cKO^* calculated by summing DEG FCs for each cortical astrocyte cluster. **(b)** Venn diagrams of DEGs found in *Arl13b^cKO^* and *Ift88^cKO^* groups (compared to Control) across cortical astrocyte Cluster 0, 1, 2. **(c)** Visualization of top DEGs in *Ift88^cKO^* cortical astrocyte clusters. **(d-f)** Top GO pathways significantly altered in *Arl13b^cKO^* and *Ift88^cKO^* cortical astrocyte clusters. **(g)** The influence of *Ift88* deletion on the key GO pathways in cortical astrocyte clusters.

**Extended Data Fig. 4.**
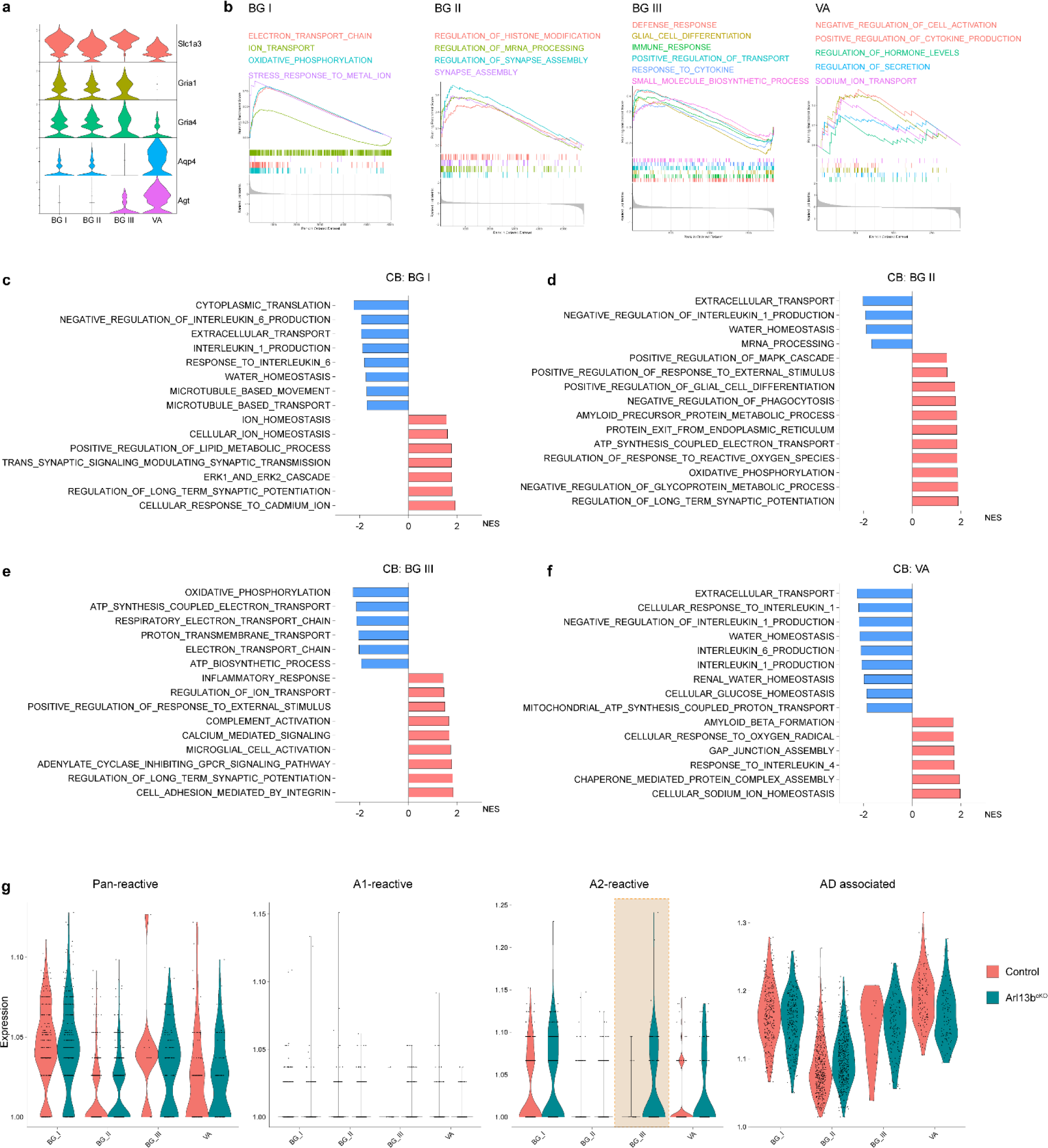
Single cell RNAseq analyses of cerebellar astrocytes. **(a)** Violin plot of the expression of genes used to distinguish Bergmann glia (BG) and velate astrocytes (VA). **(b)** GSEA plot of enriched key GO pathways in cerebellar astrocyte clusters. **(c-f)** Top GO pathways altered in *Arl13b^cKO^* cerebellar astrocyte clusters. **(g)** Gene signature expression of pan-reactive, A1-reactive, A2-reactive and Alzheimer’s disease (AD)-associated astrocytes in control and *Arl13b^cKO^* cerebellar astrocyte clusters BG_I, BG_II, BG_III, and VA. A2 reactive astrocyte signature is elevated in *Arl13b^cKO^* BG_III cluster.

**Extended Data Fig. 5.**
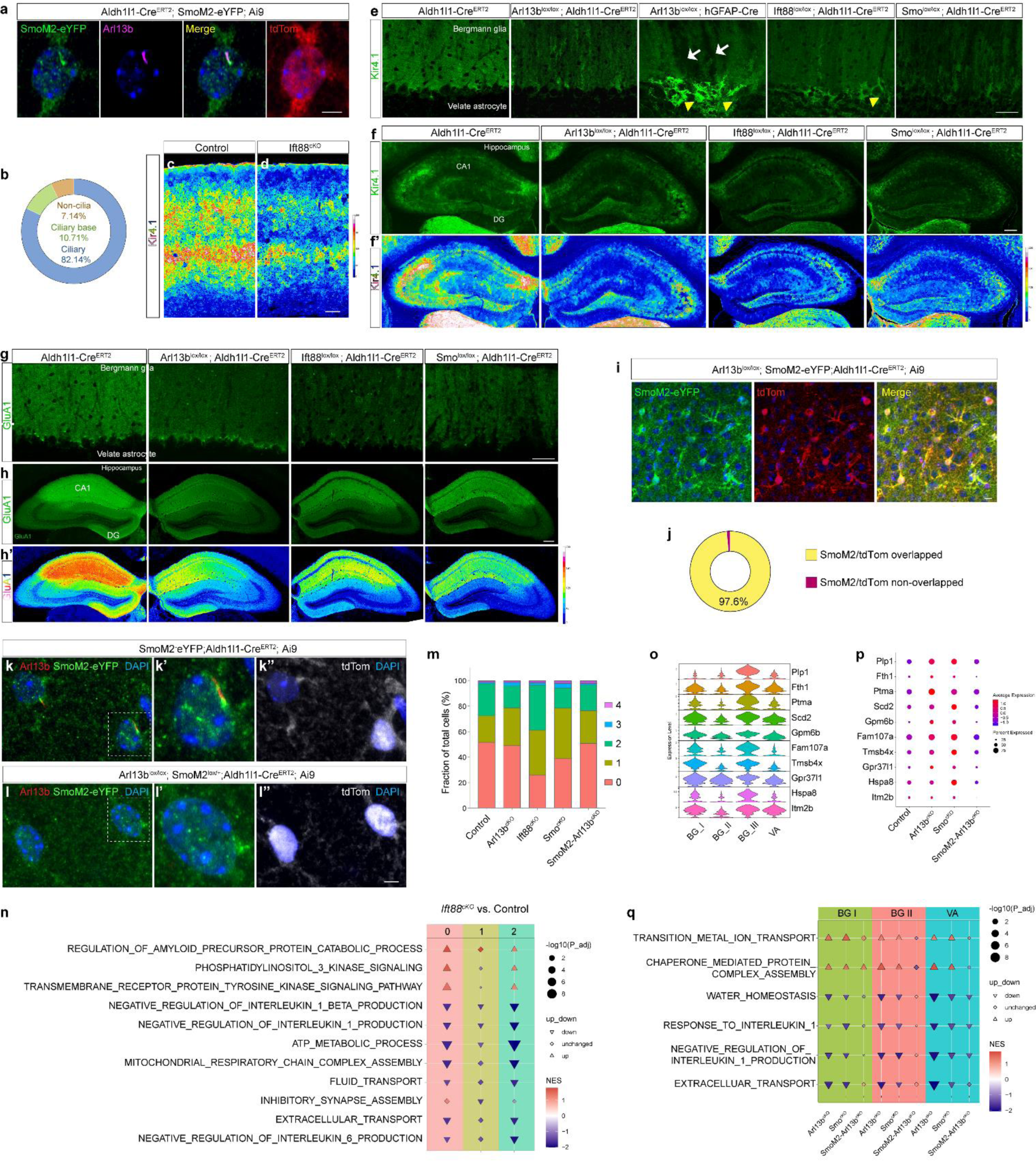
Shh signaling related changes in *Arl13b^cKO^*, *Ift88^cKO^*, and *Smo^cKO^* groups. **(a, b)** SmoM2 ciliary localization in astrocytes in *SmoM2-eYFP; Aldh1l1-Cre^ERT2^; Ai9* mice. 82.14% of SmoM2^+^ astrocytes showed SmoM2 primary ciliary localization. N=36 cells from 3 mice. **(c, d)** Kir4.1 expression patterns in Control and *Ift88^cKO^* mouse cortex. Kir4.1 expression is pseudocolored based on intensity. **(e)** Kir4.1 expression patterns in *Aldh1l1-Cre^ERT2^*, *Arl13b^lox/lox^; Aldh1l1-Cre^ERT2^*, *Arl13b^lox/lox^; hGFAP-Cre*, *Ift88^lox/lox^; Aldh1l1-Cre^ERT2^*, and *Smo^lox/lox^; Aldh1l1-Cre^ERT2^* mouse cerebellum. **(f)** Kir4.1 expression patterns in *Aldh1l1-Cre^ERT2^*, *Arl13b^lox/lox^; Aldh1l1-Cre^ERT2^*, *Ift88^lox/lox^; Aldh1l1-Cre^ERT2^*, and *Smo^lox/lox^; Aldh1l1-Cre^ERT2^* mouse hippocampus. **(f’)** Kir4.1 expression pseudocolored based on intensity in (f). **(g)** GluA1 expression patterns in *Aldh1l1-Cre^ERT2^*, *Arl13b^lox/lox^; Aldh1l1-Cre^ERT2^*, *Ift88^lox/lox^; Aldh1l1-Cre^ERT2^*, and *Smo^lox/lox^; Aldh1l1-Cre^ERT2^* mouse cerebellum. **(h)** GluA1 expression patterns in *Aldh1l1-Cre^ERT2^*, *Arl13b^lox/lox^; Aldh1l1-Cre^ERT2^*, *Ift88^lox/lox^; Aldh1l1-Cre^ERT2^*, and *Smo^lox/lox^; Aldh1l1-Cre^ERT2^* mouse hippocampus. **(h’)** GluA1 expression pseudocolored based on intensity in (h). All brains were collected at the age of 2 months. **(I, j)** Cre recombination efficiency in *Arl13b^lox/lox^; SmoM2-eYFP; Aldh1l1-Cre^ERT2^; Ai9* mice. 97.6% of all Cre^+^ (tdTom^+^) astrocytes expressed SmoM2-eYFP. All mice were administrated with tamoxifen from P7 to P9 and brains were collected at P35. **(k, l)** SmoM2-eYFP expressing astrocytes show complete loss of Arl13b protein in *SmoM2-Arl13b^cKO^* astrocytes. SmoM2-eYFP was labeled by anti-GFP antibody. **(m)** Cell percentage of cortical astrocyte clusters in all samples. **(n)** Top Cilia-cShh GO biological processes in *Arl13b^cKO^* were similarly changed in *Ift88^cKO^* cortical astrocyte clusters. **(o)** Expression of representative BGIII marker genes in cerebellar astrocytes. **(p)** Bubble plots reflecting the expression of BGIII marker genes across sample groups. **(q)** Visualization of the Cilia-cShh related GO terms that are commonly up- or down-regulated in BG and VA in *Arl13b^cKO^* and *Smo^cKO^* groups. Scale bars, 5 µm (a, i, k, l), 50 µm (c-h).

**Extended Data Fig. 6.**
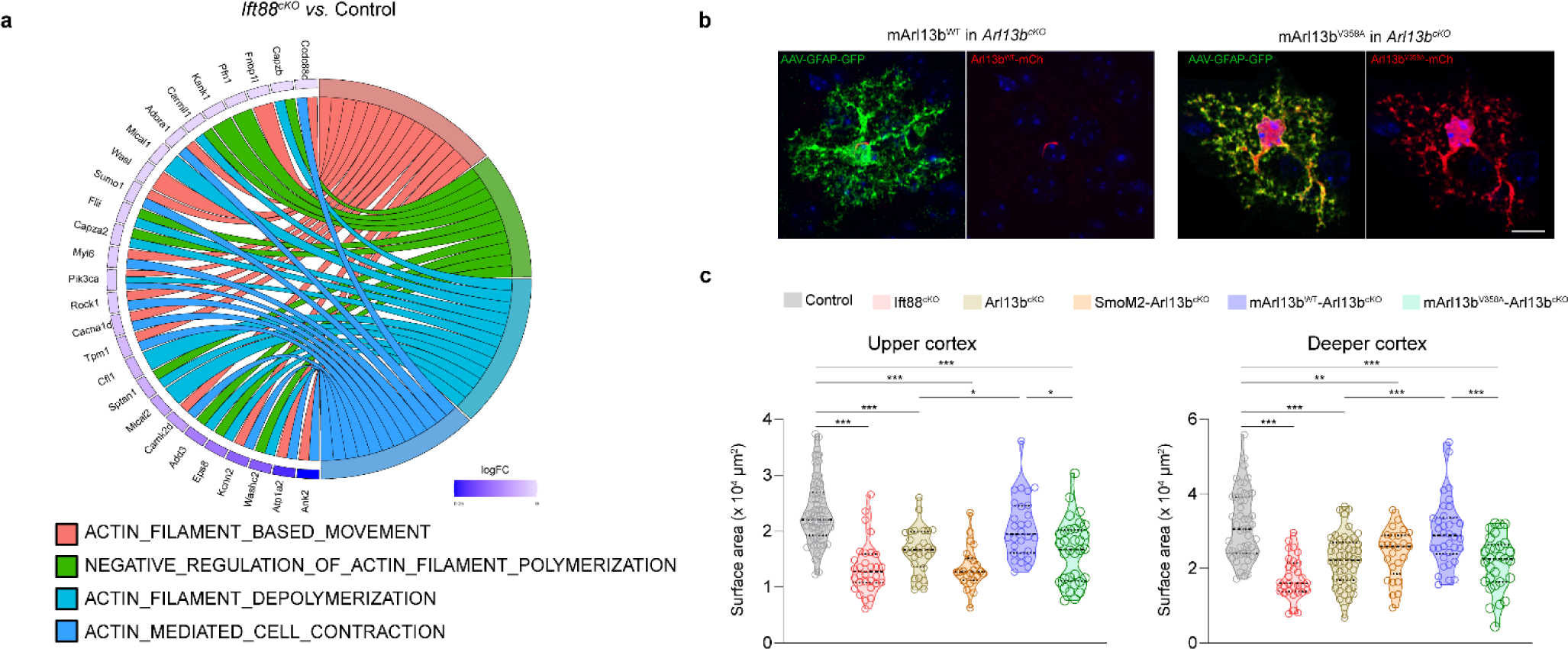
Reduced actin dynamics and morphological defects in cilia mutant astrocytes. **(a)** Circos plot of top GO biological processes enriched for actin dynamics regulation in *Ift88^cKO^* cortical Cluster 1. **(b)** The expression of mArl13b^WT^ and mArl13b^V358A^ in *Arl13b^cKO^* astrocytes. AAV viruses AAV5-GAG-DIO-mArl13b^WT^-mCh or AAV5-CAG-DIO-mArl13b^V358A^-mCh, together with AAV5-GFAP-GFP was injected in the lateral ventricles of *Arl13b^lox/lox^; Aldh1l1-Cre^ERT2^* (*Arl13b^cKO^*) brains at P1. Tamoxifen was administrated from P7 to P9. Brain sections were collected at P35. Scale bar, 5 µm. **(c)** Morphological defects in ciliary mutants. Surface area of astrocytes was measured after morphology reconstruction using IMARIS. N= 25-49 astrocytes from 3 mice per group. Comparisons of groups were performed using one-way ANOVA followed by Tukey’s test. \**P* < 0.05, \*\**P* < 0.01, \*\*\**P* < 0.001.

**Extended Data Fig.7.**
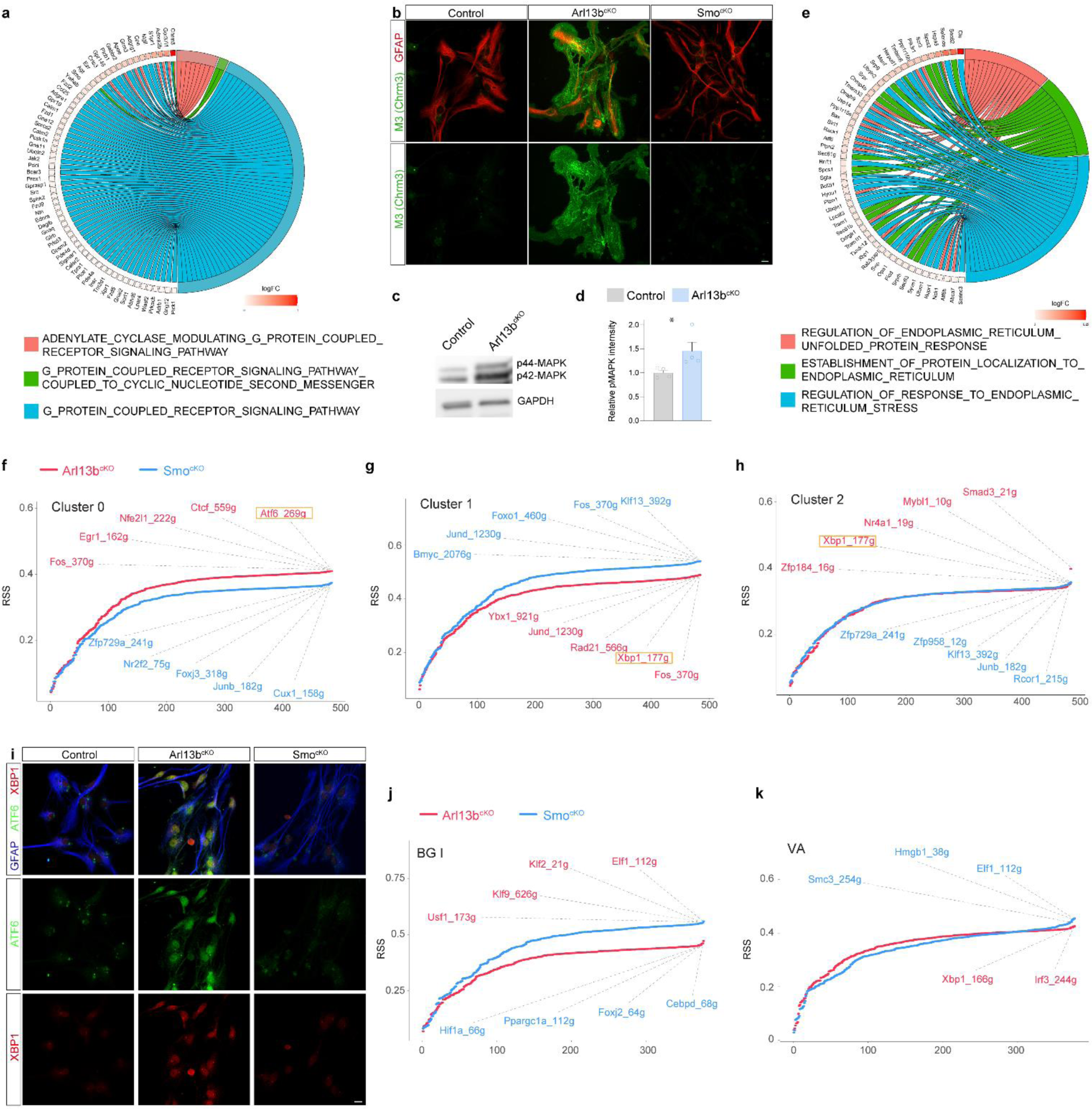
*Arl13b^cKO^* but not *Smo^cKO^* astrocytes show increased GPCR signaling activity, elevated ER stress response and distinct gene regulatory networks. **(a)** Representative circos plot of top GO biological processes enriched for GPCR signaling in *Arl13b^cKO^* cortical astrocyte cluster 1. **(b)** Expression patterns of M3 (Chrm3) in cultured control, *Arl13b^cKO^*, and *Smo^cKO^* cortical astrocytes. **(c, d)** Western blotting of phospho-MAPK using cultured control and *Arl13b^cKO^* astrocytes. N= 3 biological replicates per group. Comparison of three groups was performed using one-way-ANOVA followed by Tukey’s test. \**P* < 0.05. **(e)** Representative circos plot of cShh independent pathways enriched for ER stress in *Arl13b^cKO^* cortical astrocyte cluster 1. **(f-h)** Rank plot of *Arl13b^cKO^* and *Smo^cKO^* top regulons ordered by regulon specificity score (RSS) in cortical astrocyte clusters. **(i)** Immunostaining of XBP1 and ATF6 in cultured control, *Arl13b^cKO^*, and *Smo^cKO^* cortical astrocytes. Scale bars, 10 µm. **(j, k)** Rank plot of *Arl13b^cKO^* and *Smo^cKO^* top regulons ordered by regulon specificity score (RSS) in cerebellar astrocyte clusters. Scale bars, 10 µm.

**Extended Data Fig.8.**
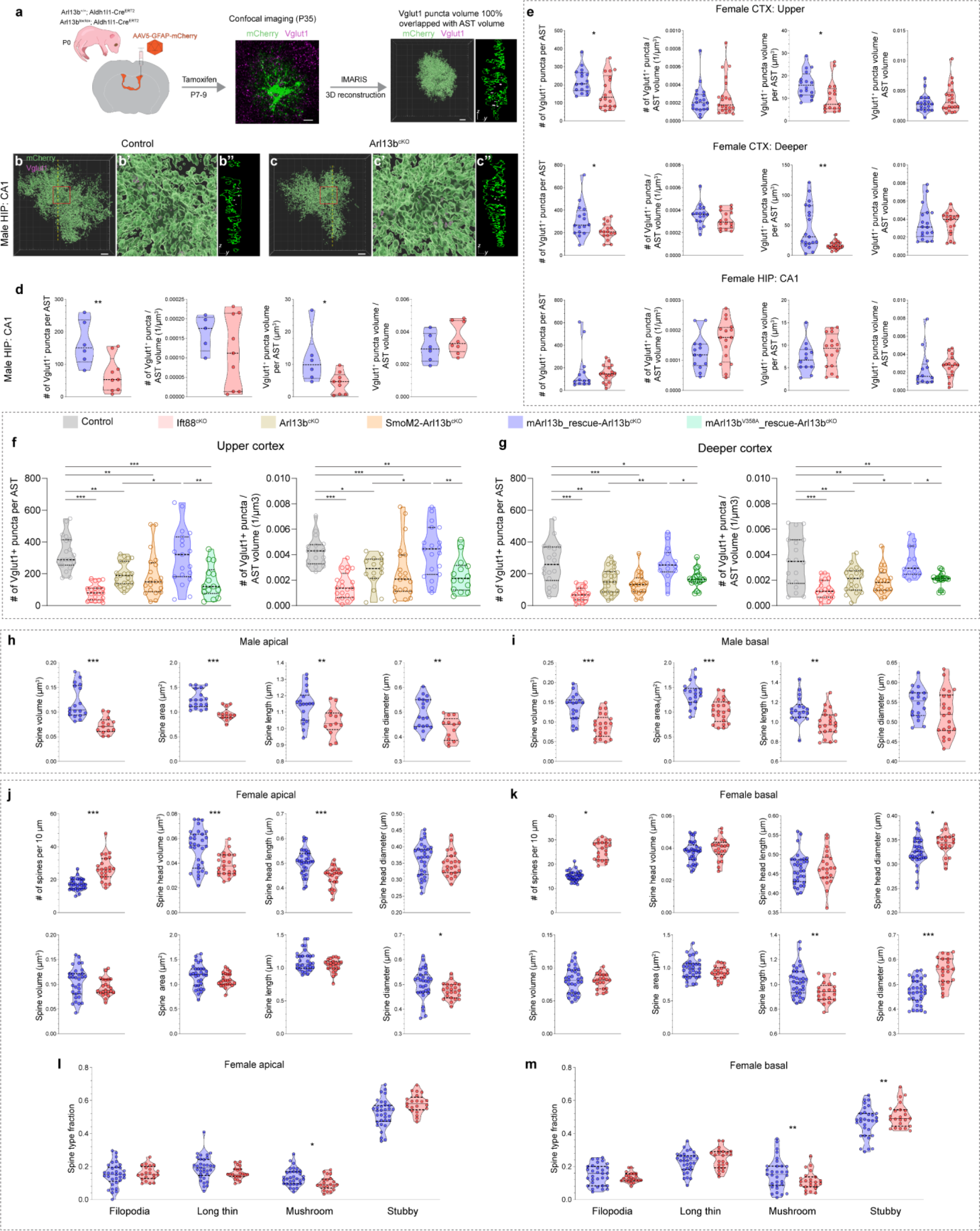
Primary ciliary dysfunction resulted in astrocytic pruning and neuronal spine defects. **(a)** Illustration of astrocytic engulfment analysis. **(b-c)** IMARIS volumetric reconstructions of astrocytes (green) containing Vglut1 (magenta) in the hippocampus in male control and *Arl13b^cKO^* brains. **(b’, c’)** Enlarged region of (b) and (c) indicated by red squares. **(b’’,c’’)** cross-section images from the yellow line indicated locations in (b) and (c). **(d)** Quantifications of the Vglut1^+^ puncta within astrocytes in the hippocampus of male mice. N= 6-9 astrocytes from 3 mice per group. **(e)** Quantifications of the Vglut1^+^ puncta within astrocytes in the upper and deeper layers of cortex, and hippocampus of female mice. N= 13-20 astrocytes from 3 mice per group. **(f, g)** Quantifications of the Vglut1^+^ puncta engulfed by upper- and deeper-cortical astrocytes in males. N=19-21 astrocytes per region from 3 mice per group. *Arl13b^cKO^* mice in the mArl13b^WT^ or mArl13b^V358A^ groups were injected with AAV5-CAG-DIO-Arl13b^WT^-mCh or AAV5-CAG-DIO-Arl13b^V358A^-mCh viruses, respectively, together with AAV5-GFAP-GFP viruses, at P1. All mice were administrated with tamoxifen from P7 to P9 and brains were collected at P35 for analysis. Vglut1^+^ puncta engulfed by astrocytes were analyzed by IMARIS. N= 25-49 astrocytes from 3 mice per group. Comparisons of six groups were performed using one-way ANOVA followed by Tukey’s test. \**P* < 0.05, \*\**P* < 0.01, \*\*\**P* < 0.001.\**P* < 0.05, \*\**P* < 0.01, \**P* < 0.001. **(h, i)** Quantifications of spine volume, area, length, diameter in apical (e) and basal (f) dendrites of male mice. N= 12-22 dendrites from three mice per region per group. **(j, k)** Quantifications of spine characteristics of M1 apical and basal dendrites from female mice. **(l, m)** Fraction of spine types in M1 cortex of female control and *Arl13b^cKO^* mice. N= 21-33 dendrites from three mice per region per group. Scale bars, 5 µm. Comparisons of two groups were performed using unpaired t-test. Comparisons of six groups were performed using one-way ANOVA followed by Tukey’s test. \**P* < 0.05, \*\**P* < 0.01, \*\*\**P* < 0.001.\**P* < 0.05, \*\**P* < 0.01, \**P* < 0.001.

**Extended Data Fig. 9.**
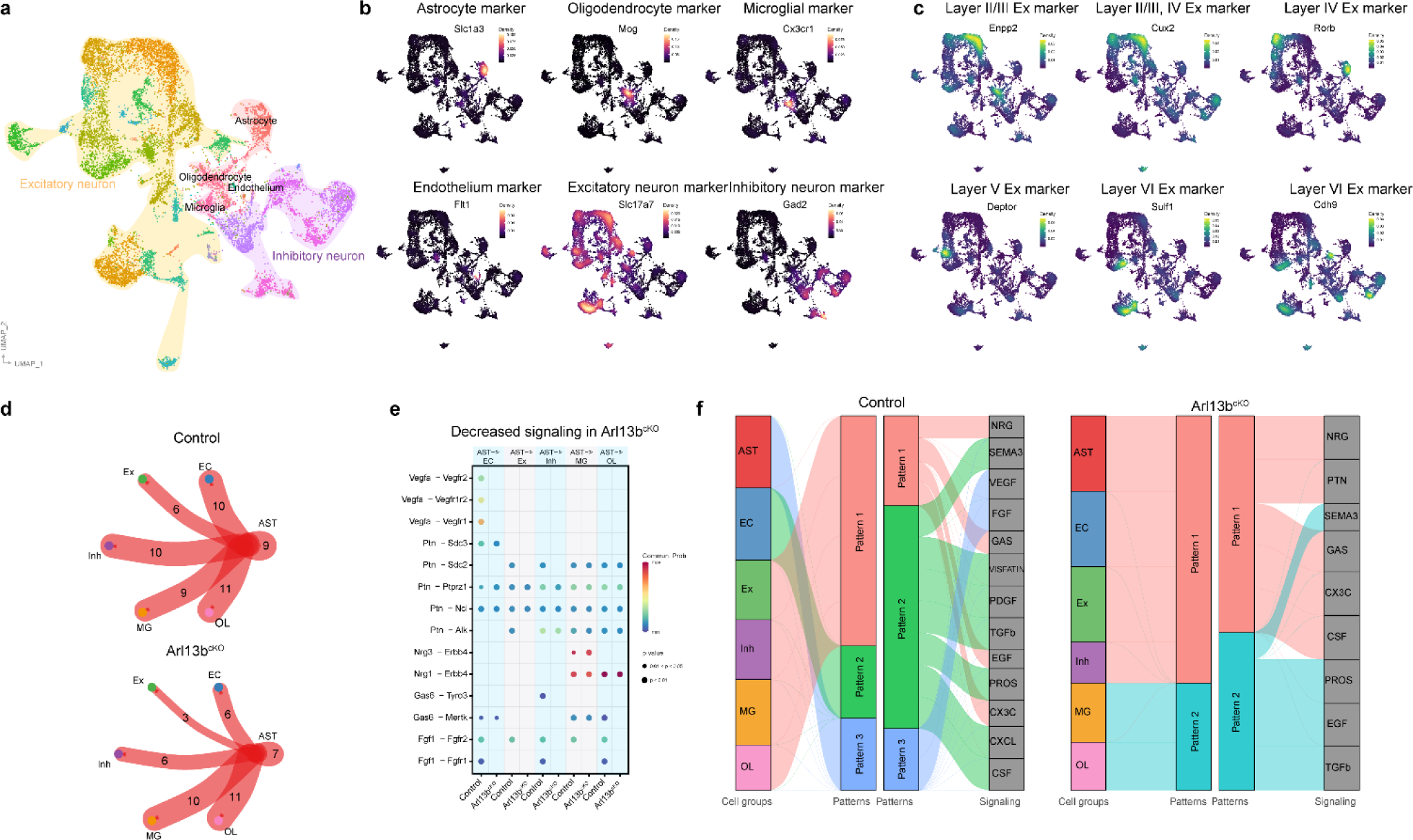
Single nucleus RNA sequencing of control and *Arl13b^cKO^* cortices. **(a)** UMAP plot of all cells from single nucleus RNA sequencing. **(b)** Density plot of representative marker genes used for cell type annotation. **(c)** Density plot of markers used to identify upper layers (Layer II/III, Layer IV) and deeper layers (Layer V, Layer VI) excitatory neurons and inhibitory neurons. **(d)** CellChat predicted number of interactions from astrocytes to other cell types in control and *Arl13b^cKO^* groups. **(e)** CellChat predicted decreased signaling from astrocytes to other cell types in *Arl13b^cKO^* compared to control brains. **(f)** CellChat predicted altered signaling interaction patterns in *Arl13b^cKO^* brains.

**Extended Data Fig. 10.**
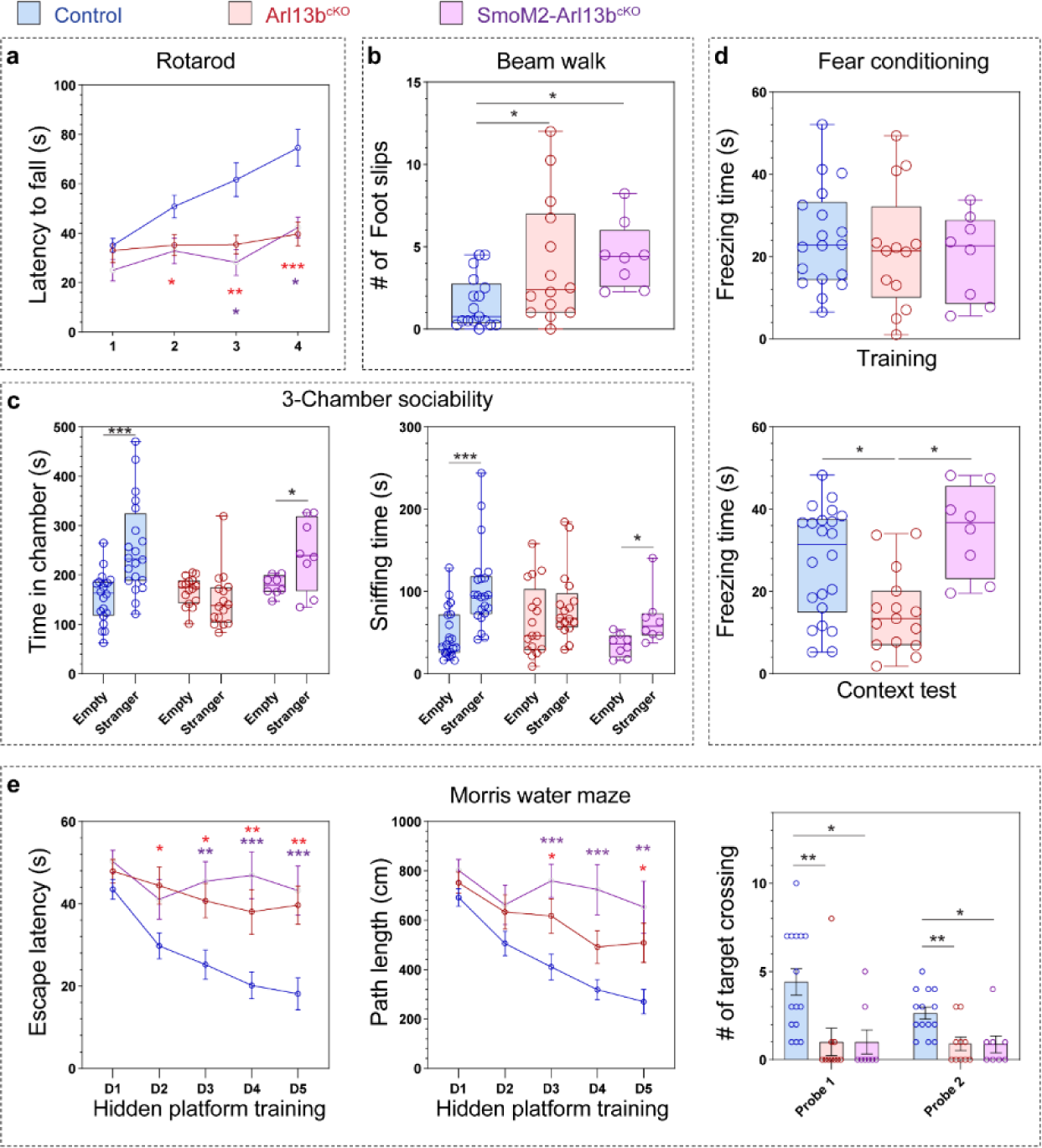
Behavior tests in control, *Arl13b^cKO^*, and *SmoM2-Arl13b^cKO^* mice. **(a)** Quantifications of the latency time to fall in the rotarod test. **(b)** Quantifications of the number of foot slips in the beam walk test. **(c)** Quantifications of the time in chamber and sniffing time in the 3-chamber sociability test. **(d)** Quantifications of % of freezing time in the fear conditioning test. **(e)** Quantifications for the Morris water maze test. N= 8-16 mice per group. Comparisons were performed using one-way ANOVA followed by Tukey’s test. * in red, *Arl13b^cKO^ vs.* Control; * in purple, *SmoM2-Arl13b^cKO^ vs.* Control. \**P* < 0.05, \*\**P* < 0.01, \*\*\**P* < 0.001.

## Notes

### Competing Interest Statement

The authors have declared no competing interest.

## References

1. Cahoy, J.D., et al. A Transcriptome Database for Astrocytes, Neurons, and Oligodendrocytes: A New Resource for Understanding Brain Development and Function. J Neurosci 28, 264–278 (2008).

2. Sloan, S.A. & Barres, B.A. Mechanisms of astrocyte development and their contributions to neurodevelopmental disorders. Curr Opin Neurobiol 27, 75–81 (2014).

3. Clarke, L.E. & Barres, B.A. Emerging roles of astrocytes in neural circuit development. Nat Rev Neurosci 14, 311–321 (2013).

4. Liddelow, S.A. & Barres, B.A. Reactive Astrocytes: Production, Function, and Therapeutic Potential. Immunity 46, 957–967 (2017).

5. V, M.A., et al. Astrocytes and disease: a neurodevelopmental perspective. Genes & development 26, 891–907.

6. Barker, A.J. & Ullian, E.M. Astrocytes and Synaptic Plasticity. Neurosci 16, 40–50 (2010).

7. Chung, W.-S., Allen, N.J. & Eroglu, C. Astrocytes Control Synapse Formation, Function, and Elimination. Csh Perspect Biol 7, a020370 (2015).

8. Ceprian, M. & Fulton, D. Glial Cell AMPA Receptors in Nervous System Health, Injury and Disease. Int J Mol Sci 20 (2019).

9. Siracusa, R., Fusco, R. & Cuzzocrea, S. Astrocytes: Role and Functions in Brain Pathologies. Front Pharmacol 10, 1114 (2019).

10. Eroglu, C. & Barres, B.A. Regulation of synaptic connectivity by glia. Nature 468, 223–231 (2010).

11. Todd, F.W., et al. Neurons diversify astrocytes in the adult brain through sonic hedgehog signaling. Science 351, 849–854.

12. Batiuk, M.Y., et al. Identification of region-specific astrocyte subtypes at single cell resolution. Nat Commun 11, 1220 (2020).

13. Bayraktar, O.A., Fuentealba, L.C., Alvarez-Buylla, A. & Rowitch, D.H. Astrocyte Development and Heterogeneity. Csh Perspect Biol 7, a020362 (2015).

14. Chai, H., et al. Neural Circuit-Specialized Astrocytes: Transcriptomic, Proteomic, Morphological, and Functional Evidence. Neuron 95, 531–549.e539 (2017).

15. Molofsky, A.V. & Deneen, B. Astrocyte development: A Guide for the Perplexed. Glia 63, 1320–1329 (2015).

16. Khakh, B.S. & Deneen, B. The Emerging Nature of Astrocyte Diversity. Annu Rev Neurosci 42, 187–207 (2019).

17. Farmer, W.T. & Murai, K. Resolving Astrocyte Heterogeneity in the CNS. Front Cell Neurosci 11, 300 (2017).

18. Gingrich, E.C., Case, K. & Garcia, A.D.R. A subpopulation of astrocyte progenitors defined by Sonic hedgehog signaling. Neural Dev 17, 2 (2022).

19. Hill, S.A., et al. Sonic hedgehog signaling in astrocytes mediates cell type-specific synaptic organization. Elife 8, e45545 (2019).

20. Xie, Y., et al. Astrocyte-neuron crosstalk through Hedgehog signaling mediates cortical synapse development. Cell Reports 38, 110416 (2022).

21. Garcia, A.D.R., Petrova, R., Eng, L. & Joyner, A.L. Sonic Hedgehog Regulates Discrete Populations of Astrocytes in the Adult Mouse Forebrain. J Neurosci 30, 13597–13608 (2010).

22. Stogsdill, J.A., et al. Astrocytic neuroligins control astrocyte morphogenesis and synaptogenesis. Nature 551, 192–197 (2017).

23. Gerald, S., Karl, S. & Christian, S. Astrocyte dysfunction in neurological disorders: a molecular perspective. Nat Rev Neurosci 7, 194–206.

24. Lee, H.-G., Wheeler, M.A. & Quintana, F.J. Function and therapeutic value of astrocytes in neurological diseases. Nat Rev Drug Discov 21, 339–358 (2022).

25. Lin, C.-C.J., et al. Identification of diverse astrocyte populations and their malignant analogs. Nat Neurosci 20, 396–405 (2017).

26. Yu, X., et al. Reducing Astrocyte Calcium Signaling In Vivo Alters Striatal Microcircuits and Causes Repetitive Behavior. Neuron 99, 1170–1187.e1179 (2018).

27. Bronzuoli, M.R., et al. Neuroglia in the autistic brain: evidence from a preclinical model. Mol Autism 9, 66 (2018).

28. Petrelli, F., Pucci, L. & Bezzi, P. Astrocytes and Microglia and Their Potential Link with Autism Spectrum Disorders. Front Cell Neurosci 10, 21 (2016).

29. Wheway, G., Nazlamova, L. & Hancock, J.T. Signaling through the Primary Cilium. Frontiers Cell Dev Biology 6, 8 (2018).

30. Schou, K.B., Pedersen, L.B. & Christensen, S.T. Ins and outs of GPCR signaling in primary cilia. EMBO reports 16, 1099–1113 (2015).

31. Christensen, S.T., Clement, C.A., Satir, P. & Pedersen, L.B. Primary cilia and coordination of receptor tyrosine kinase (RTK) signalling. The Journal of Pathology 226, 172–184 (2012).

32. Hilgendorf, K.I., Johnson, C.T. & Jackson, P.K. The primary cilium as a cellular receiver: organizing ciliary GPCR signaling. Curr Opin Cell Biol 39, 84–92 (2016).

33. Delling, M., DeCaen, P.G., Doerner, J.F., Febvay, S. & Clapham, D.E. Primary cilia are specialized calcium signalling organelles. Nature 504, 311–314 (2013).

34. Guo, J., et al. Primary Cilia Signaling Shapes the Development of Interneuronal Connectivity. DEVCEL 42, 286–300.e284 (2017).

35. Phua, S.C., Lin, Y.-C. & Inoue, T. An intelligent nano-antenna: Primary cilium harnesses TRP channels to decode polymodal stimuli. Cell Calcium 58, 415–422 (2015).

36. Badano, J.L., Mitsuma, N., Beales, P.L. & Katsanis, N. The ciliopathies: an emerging class of human genetic disorders. Annual Review of Genomics and Human Genetics 7, 125–148 (2006).

37. Hildebrandt, F., of, T.B.N.E.J. & 2011. Ciliopathies. Mass Medical Soc 364, 1533–1543 (2011).

38. Novarino, G., Akizu, N. & Gleeson, J.G. Modeling Human Disease in Humans: The Ciliopathies. Cell 147, 70–79 (2011).

39. Wheway, G. & Mitchison, H.M. Opportunities and Challenges for Molecular Understanding of Ciliopathies-The 100,000 Genomes Project. Frontiers in genetics 10, 127 (2019).

40. Guo, J., et al. Developmental disruptions underlying brain abnormalities in ciliopathies. Nature communications 6, 1–13 (2015).

41. Higginbotham, H., et al. Arl13b-regulated cilia activities are essential for polarized radial glial scaffold formation. Nature neuroscience, 1–9 (2013).

42. Guo, J., et al. Primary Cilia Signaling Promotes Axonal Tract Development and Is Disrupted in Joubert Syndrome-Related Disorders Models. Developmental Cell 51, 759–774.e755 (2019).

43. Alvarez Retuerto, A.I., et al. Association of common variants in the Joubert syndrome gene (AHI1) with autism. Human Molecular Genetics 17, 3887–3896 (2008).

44. Brancati, F., Dallapiccola, B. & Valente, E. Joubert Syndrome and related disorders. Orphanet Journal of Rare Diseases 5, 20–10 (2010).

45. Cantagrel, V., et al. Mutations in the Cilia Gene ARL13B Lead to the Classical Form of Joubert Syndrome. The American Journal of Human Genetics 83, 170–179 (2008).

46. Iskender, C.T., Tarim, E. & Alkan, O. Joubert syndrome and related disorders, prenatal diagnosis with ultrasound and magnetic resonance imaging. Journal of the Turkish German Gynecological Association, 1–4 (2011).

47. Poretti, A., et al. Diffusion Tensor Imaging in Joubert Syndrome. American Journal of Neuroradiology 28, 1929–1933 (2007).

48. Senocak, E.U., Oguz, K.K., Haliloglu, G., Topçu, M. & Cila, A. Structural abnormalities of the brain other than molar tooth sign in joubert syndrome related disorders. Diagnostic and Interventional Radiology, 1–4 (2009).

49. Rafiullah, R., et al. A novel homozygous ARL13B variant in patients with Joubert syndrome impairs its guanine nucleotide-exchange factor activity. European Journal of Human Genetics 25, 1324–1334 (2017).

50. Migliavacca, E., et al. A Potential Contributory Role for Ciliary Dysfunction in the 16p11.2 600 kb BP4-BP5 Pathology. American journal of human genetics 96, 784–796 (2015).

51. Alicia, G.-G., G, C.N. & G, G.J. Primary Cilia in the Developing and Mature Brain. Neuron 82, 511–521.

52. A, G.J. & Kirk, M. Neuronal ciliary signaling in homeostasis and disease. Cell Mol Life Sci 67, 3287–3297.

53. A, A.-A., et al. Primary Cilia Regulate Proliferation of Amplifying Progenitors in Adult Hippocampus: Implications for Learning and Memory. J Neurosci 31, 9933–9944.

54. Min, P.S., Jin, J.H. & Ho, L.J. Roles of Primary Cilia in the Developing Brain. Front Cell Neurosci 13, 1358.

55. Sheu, S.-H., et al. A serotonergic axon-cilium synapse drives nuclear signaling to maintain chromatin accessibility. Biorxiv, 2021.2009.2027.461878 (2021).

56. Jiami, G., et al. Primary Cilia Signaling Promotes Axonal Tract Development and Is Disrupted in Joubert Syndrome-Related Disorders Models. Dev Cell 51, 759–774.e755.

57. M, G.S., et al. Arborization of Dendrites by Developing Neocortical Neurons Is Dependent on Primary Cilia and Type 3 Adenylyl Cyclase. J Neurosci 33, 2626–2638.

58. Kang, K. & Song, M.-R. Diverse FGF receptor signaling controls astrocyte specification and proliferation. Biochem Bioph Res Co 395, 324–329 (2010).

59. Savchenko, E., et al. FGF family members differentially regulate maturation and proliferation of stem cell-derived astrocytes. Sci Rep-uk 9, 9610 (2019).

60. Estefania, A.-F., Ana, O.-R., Iñigo, A., M, G.-S.L. & Maria-Angeles, A. Notch signaling in astrocytes mediates their morphological response to an inflammatory challenge. Cell Death Discov 5, 85–14 (2019).

61. Su, C.Y., Bay, S.N., Mariani, L.E., Hillman, M.J. & Caspary, T. Temporal deletion of Arl13b reveals that a mispatterned neural tube corrects cell fate over time. Development 139, 4062–4071 (2012).

62. Haycraft, C.J., et al. Intraflagellar transport is essential for endochondral bone formation. Development 134, 307–316 (2007).

63. Koike, K., et al. Danger perception and stress response through an olfactory sensor for the bacterial metabolite hydrogen sulfide. Neuron 109, 2469–2484 e2467 (2021).

64. Humbert, M.C., et al. ARL13B, PDE6D, and CEP164 form a functional network for INPP5E ciliary targeting. Proceedings of the National Academy of Sciences of the United States of America 109, 19691–19696 (2012).

65. Pazour, G.J., et al. Chlamydomonas IFT88 and its mouse homologue, polycystic kidney disease gene tg737, are required for assembly of cilia and flagella. J Cell Biol 151, 709–718 (2000).

66. Freeman, M.R. Specification and morphogenesis of astrocytes. Science 330, 774–778 (2010).

67. Forcioli-Conti, N., Esteve, D., Bouloumie, A., Dani, C. & Peraldi, P. The size of the primary cilium and acetylated tubulin are modulated during adipocyte differentiation: Analysis of HDAC6 functions in these processes. Biochimie 124, 112–123 (2016).

68. Caspary, T., Larkins, C.E. & Anderson, K.V. The graded response to Sonic Hedgehog depends on cilia architecture. Dev Cell 12, 767–778 (2007).

69. Matusova, Z., Hol, E.M., Pekny, M., Kubista, M. & Valihrach, L. Reactive astrogliosis in the era of single-cell transcriptomics. Front Cell Neurosci 17, 1173200 (2023).

70. Hill, S.A., Fu, M. & Garcia, A.D.R. Sonic hedgehog signaling in astrocytes. Cell Mol Life Sci 78, 1393–1403 (2021).

71. Tamara, C., E, L.C. & V, A.K. The Graded Response to Sonic Hedgehog Depends on Cilia Architecture. Developmental Cell 12, 767–778.

72. Guemez-Gamboa, A., Coufal, N.G. & Gleeson, J.G. Primary Cilia in the Developing and Mature Brain. Neuron 82, 511–521 (2014).

73. Bangs, F. & Anderson, K.V. Primary Cilia and Mammalian Hedgehog Signaling. Csh Perspect Biol 9, a028175 (2017).

74. Corbit, K.C., et al. Vertebrate Smoothened functions at the primary cilium. Nature 437, 1018–1021 (2005).

75. Xie, J., et al. Activating Smoothened mutations in sporadic basal-cell carcinoma. Nature 391, 90–92 (1998).

76. Garcia, A.D.R., et al. The Elegance of Sonic Hedgehog: Emerging Novel Functions for a Classic Morphogen. J Neurosci 38, 9338–9345 (2018).

77. Zhuo, L., et al. hGFAP-cre transgenic mice for manipulation of glial and neuronal function in vivo. Genesis 31, 85–94 (2001).

78. Szu, J.I. & Binder, D.K. The Role of Astrocytic Aquaporin-4 in Synaptic Plasticity and Learning and Memory. Frontiers Integr Neurosci 10, 8 (2016).

79. Siehler, S. Regulation of RhoGEF proteins by G12/13-coupled receptors. Brit J Pharmacol 158, 41–49 (2009).

80. Higginbotham, H., et al. Arl13b in primary cilia regulates the migration and placement of interneurons in the developing cerebral cortex. Dev Cell 23, 925–938 (2012).

81. Gigante, E.D., Taylor, M.R., Ivanova, A.A., Kahn, R.A. & Caspary, T. ARL13B regulates Sonic hedgehog signaling from outside primary cilia. Elife 9 (2020).

82. Mariani, L.E., et al. Arl13b regulates Shh signaling from both inside and outside the cilium. Mol Biol Cell 27, 3780–3790 (2016).

83. Ferent, J., et al. The Ciliary Protein Arl13b Functions Outside of the Primary Cilium in Shh-Mediated Axon Guidance. Cell Rep 29, 3356–3366 e3353 (2019).

84. Nakagawa, N., et al. Memo1-Mediated Tiling of Radial Glial Cells Facilitates Cerebral Cortical Development. Neuron 103, 836–852.e835 (2019).

85. Gutkind, J.S. The Pathways Connecting G Protein-coupled Receptors to the Nucleus through Divergent Mitogen-activated Protein Kinase Cascades*. J Biol Chem 273, 1839–1842 (1998).

86. Aibar, S., et al. SCENIC: single-cell regulatory network inference and clustering. Nat Methods 14, 1083–1086 (2017).

87. Sprenkle, N.T., Sims, S.G., Sánchez, C.L. & Meares, G.P. Endoplasmic reticulum stress and inflammation in the central nervous system. Mol Neurodegener 12, 42 (2017).

88. Endo, F., et al. Molecular basis of astrocyte diversity and morphology across the CNS in health and disease. Science 378, eadc9020 (2022).

89. Freeman, M.R. Specification and Morphogenesis of Astrocytes. Science 330, 774–778 (2010).

90. Schiweck, J., Eickholt, B.J. & Murk, K. Important Shapeshifter: Mechanisms Allowing Astrocytes to Respond to the Changing Nervous System During Development, Injury and Disease. Front Cell Neurosci 12, 261 (2018).

91. Lawal, O., Severino, F.P.U. & Eroglu, C. The role of astrocyte structural plasticity in regulating neural circuit function and behavior. Glia 70, 1467–1483 (2022).

92. Feng, G., et al. Imaging Neuronal Subsets in Transgenic Mice Expressing Multiple Spectral Variants of GFP. Neuron 28, 41–51 (2000).

93. Jin, S., et al. Inference and analysis of cell-cell communication using CellChat. Nat Commun 12, 1088 (2021).

94. Thomas, S., et al. Identification of a novel ARL13B variant in a Joubert syndrome-affected patient with retinal impairment and obesity. Eur J Hum Genet 23, 621–627 (2015).

95. Cantagrel, V., et al. Mutations in the cilia gene ARL13B lead to the classical form of Joubert syndrome. Am J Hum Genet 83, 170–179 (2008).

96. Juric-Sekhar, G., Adkins, J., Doherty, D. & Hevner, R.F. Joubert syndrome: brain and spinal cord malformations in genotyped cases and implications for neurodevelopmental functions of primary cilia. Acta Neuropathol 123, 695–709 (2012).

97. Miertzschke, M., Koerner, C., Spoerner, M. & Wittinghofer, A. Structural insights into the small G-protein Arl13B and implications for Joubert syndrome. Biochem J 457, 301–311 (2014).

98. Parisi, M.A. The molecular genetics of Joubert syndrome and related ciliopathies: The challenges of genetic and phenotypic heterogeneity. Transl Sci Rare Dis 4, 25–49 (2019).

99. Rafiullah, R., et al. A novel homozygous ARL13B variant in patients with Joubert syndrome impairs its guanine nucleotide-exchange factor activity. Eur J Hum Genet 25, 1324–1334 (2017).

100. Chung, W.-S., et al. Astrocytes mediate synapse elimination through MEGF10 and MERTK pathways. Nature 504, 394–400 (2013).

101. Morizawa, Y.M., et al. Synaptic pruning through glial synapse engulfment upon motor learning. Nat Neurosci 25, 1458–1469 (2022).

102. Russo, F.B., et al. Modeling the Interplay Between Neurons and Astrocytes in Autism Using Human Induced Pluripotent Stem Cells. Biol Psychiat 83, 569–578 (2018).

103. Jiami, G., et al. Developmental disruptions underlying brain abnormalities in ciliopathies. Nat Commun 6, 1–13.

104. Wang, Y., et al. Melanocortin 4 receptor signals at the neuronal primary cilium to control food intake and body weight. J Clin Invest 131, e142064 (2021).

105. Higginbotham, H., et al. Arl13b in primary cilia regulates the migration and placement of interneurons in the developing cerebral cortex. Developmental Cell 23, 925–938 (2012).

106. DeMars, K.M., Ross, M.R., Starr, A. & McIntyre, J.C. Neuronal primary cilia integrate peripheral signals with metabolic drives. Front Physiol 14, 1150232 (2023).

107. Huang, S., Xing, Y. & Liu, Y. Emerging roles for the ER stress sensor IRE1α in metabolic regulation and disease. J Biol Chem 294, 18726–18741 (2019).

108. A, L.M. & G, G.J. The primary cilium as a cellular signaling center: lessons from disease. Curr Opin Genet Dev 19, 220–229.

109. F, H. & Journal, o.T.B.N.E. Ciliopathies. New Engl J Med 364, 1533–1543.

110. U, M.D., B, R.R., D, L.R. & Developmental, c.C.M.A. Proteomics of primary cilia by proximity labeling. Elsevier 35, 497–512 (2015).

111. Aslanyan, M.G., et al. A targeted multi-proteomics approach generates a blueprint of the ciliary ubiquitinome. Frontiers Cell Dev Biology 11, 1113656 (2023).

112. Sigg, M.A., et al. Evolutionary Proteomics Uncovers Ancient Associations of Cilia with Signaling Pathways. Dev Cell 43, 744–762.e711 (2017).

113. Ma, R., Kutchy, N.A., Chen, L., Meigs, D.D. & Hu, G. Primary cilia and ciliary signaling pathways in aging and age-related brain disorders. Neurobiol Dis 163, 105607 (2021).

114. F, B.N., et al. Hippocampal and Cortical Primary Cilia Are Required for Aversive Memory in Mice. Plos One 9, e106576–106510.

115. Karunakaran, K.B., Chaparala, S., Lo, C.W. & Ganapathiraju, M.K. Cilia interactome with predicted protein–protein interactions reveals connections to Alzheimer’s disease, aging and other neuropsychiatric processes. Sci Rep-uk 10, 15629 (2020).

116. Schmidt, S., et al. Primary cilia and SHH signaling impairments in human and mouse models of Parkinson’s disease. Nat Commun 13, 4819 (2022).

117. Sofroniew, M.V. Astrocyte Reactivity: Subtypes, States, and Functions in CNS Innate Immunity. Trends Immunol 41, 758–770 (2020).

118. Bouvier, D.S., et al. The Multifaceted Neurotoxicity of Astrocytes in Ageing and Age-Related Neurodegenerative Diseases: A Translational Perspective. Front Physiol 13, 814889 (2022).

119. Andrzej, C.A. & Zbigniew, S. Bergmann Glia, Long-Term Depression, and Autism Spectrum Disorder. Mol Neurobiol 54, 1156–1166.

120. Ajoolabady, A., Lindholm, D., Ren, J. & Pratico, D. ER stress and UPR in Alzheimer’s disease: mechanisms, pathogenesis, treatments. Cell Death Dis 13, 706 (2022).

121. Long, F., Zhang, X.M., Karp, S., Yang, Y. & McMahon, A.P. Genetic manipulation of hedgehog signaling in the endochondral skeleton reveals a direct role in the regulation of chondrocyte proliferation. Development 128, 5099–5108 (2001).

122. Ganat, Y.M., et al. Early postnatal astroglial cells produce multilineage precursors and neural stem cells in vivo. J Neurosci 26, 8609–8621 (2006).

123. Madisen, L., et al. A robust and high-throughput Cre reporting and characterization system for the whole mouse brain. Nat Neurosci 13, 133–140 (2010).

124. Nakagawa, N., et al. Memo1-Mediated Tiling of Radial Glial Cells Facilitates Cerebral Cortical Development. Neuron 103, 836–852 e835 (2019).

125. Jeong, J., Mao, J., Tenzen, T., Kottmann, A.H. & McMahon, A.P. Hedgehog signaling in the neural crest cells regulates the patterning and growth of facial primordia. Genes Dev 18, 937–951 (2004).

126. Zhang, Y., et al. Purification and Characterization of Progenitor and Mature Human Astrocytes Reveals Transcriptional and Functional Differences with Mouse. Neuron 89, 37–53 (2016).

127. Chen, J.K., Taipale, J., Cooper, M.K. & Beachy, P.A. Inhibition of Hedgehog signaling by direct binding of cyclopamine to Smoothened. Genes Dev 16, 2743–2748 (2002).

128. Byun, Y.G. & Chung, W.S. A Novel In Vitro Live-imaging Assay of Astrocyte-mediated Phagocytosis Using pH Indicator-conjugated Synaptosomes. J Vis Exp (2018).

129. Auguste, Y.S.S., et al. Oligodendrocyte precursor cells engulf synapses during circuit remodeling in mice. Nat Neurosci 25, 1273–1278 (2022).

130. Zheng, G.X., et al. Massively parallel digital transcriptional profiling of single cells. Nat Commun 8, 14049 (2017).

131. Hafemeister, C. & Satija, R. Normalization and variance stabilization of single-cell RNA-seq data using regularized negative binomial regression. Genome Biol 20, 296 (2019).

132. Korsunsky, I., et al. Fast, sensitive and accurate integration of single-cell data with Harmony. Nat Methods 16, 1289–1296 (2019).

133. Wilk, A.J., et al. Multi-omic profiling reveals widespread dysregulation of innate immunity and hematopoiesis in COVID-19. J Exp Med 218 (2021).

134. Alquicira-Hernandez, J. & Powell, J.E. Nebulosa recovers single-cell gene expression signals by kernel density estimation. Bioinformatics 37, 2485–2487 (2021).

135. Aibar, S., et al. SCENIC: single-cell regulatory network inference and clustering. Nat Methods 14, 1083–1086 (2017).

136. Jin, S., et al. Inference and analysis of cell-cell communication using CellChat. Nat Commun 12, 1088 (2021).

137. Vorhees, C.V. & Williams, M.T. Morris water maze: procedures for assessing spatial and related forms of learning and memory. Nat Protoc 1, 848–858 (2006).

